# Hippocampal hub neurons maintain unique functional properties throughout their lifetime

**DOI:** 10.1101/813923

**Authors:** Marco Bocchio, Claire Gouny, David Angulo-Garcia, Tom Toulat, Thomas Tressard, Agnès Baude, Rosa Cossart

**Affiliations:** INMED (INSERM U1249), Aix-Marseille University, Turing Center for Living Systems, Parc Scientifique de Luminy, 13273, Marseille, France; Grupo de Modelado Computacional–Dinámica y Complejidad de Sistemas, Instituto de Matemáticas Aplicadas, Universidad de Cartagena, 130001, Cartagena, Colombia

## Abstract

The temporal embryonic origins of cortical GABA neurons are critical for their specialization. In the neonatal hippocampus, GABA cells born the earliest (ebGABAs) operate as ‘hubs’ by orchestrating neuronal dynamics. However, their fate remains largely unknown. To fill this gap, we have examined CA1 ebGABAs using a combination of electrophysiology, neurochemical analysis, optogenetic connectivity mapping as well as *ex vivo* and *in vivo* calcium imaging. We show that CA1 ebGABAs not only operate as hubs during development, but also maintain distinct morpho-physiological and connectivity profiles, including a bias for long-range targets and local excitatory inputs. *In vivo*, ebGABAs signal a variety of network states, including the activation of local CA1 assemblies. Hence, ebGABAs are specified from birth to ensure unique functions throughout their lifetime. In the adult brain, this may take the form of a long-range hub role through the coordination of cell assemblies across distant regions.

## Introduction

GABAergic neurons are a critical component of cortical circuit development and function. This sparse and heterogeneous population of cells has been classified according to several parameters, including genetic and molecular markers, connectivity schemes as well as morphological and electrophysiological properties^1^. According to leading theories, this heterogeneity greatly enhances the computational power of cortical networks ^2^.

The diversity among GABAergic cells is specified as early as progenitor stages in the ganglionic eminences, the embryonic regions that give rise to cortical inhibitory neurons^3, 4^. Genetic restriction of neuronal potential from spatially distributed progenitors is a major determinant of GABA neuron diversity^5–7^. However, a temporal clock also shapes GABA neuron fate: discrete temporal windows within the same ganglionic eminence specify different cell types^8, 9^. Hence, place and time of origin from discrete progenitor pools in the ganglionic eminences determine, at least in part, the ultimate features that the cell will display in adult cortical circuits.

This distinct temporal control of GABA neurons’ fate is particularly striking when considering early-born cells. In the CA3 region of the hippocampus, early-born GABAergic neurons (ebGABAs) give rise to cells acting as ‘hubs’ during the perinatal period, i.e. combining exceptional functional and effective connectivity degrees^10^. Due to this remarkable connectivity scheme, CA3 hub cells anticipate and coordinate single-handedly spontaneous network bursts occurring in the form of giant depolarizing potentials (GDPs) in postnatal slices^10, 11^. Interestingly, these cells are still present in adult animals and a fraction of them display long-range projections to the medial septum^12^.

It is not known whether a single ebGABA is able to orchestrate network synchrony in the entire hippocampal formation or only in CA3, a region that is characterized by notable recurrent connectivity and that is the preferred site of GDP initiation^13^. In addition, despite their major developmental role, the integration and function of ebGABAs into adult circuits remains unknown. More generally, it is not known whether an early neuronal birth date leads to distinct functional properties in adult circuits. To fill this gap, we have examined ebGABAs in the CA1 region of the hippocampus, because this area is well understood in terms of cell types and connectivity^14^ and is more accessible for *in vivo* calcium imaging. We investigated ebGABAs’ morphological and electrophysiological properties, their local and long-range connectivity as well as their involvement in network dynamics. We show that CA1 ebGABAs operate as hub cells during the early postnatal period and maintain distinct properties in adulthood, encompassing neurochemical content, intrinsic firing patterns, input connectivity and *in vivo* activity.

## Results

### EbGABAs are operational hub cells in the developing CA1

We sought to test whether ebGABAs played a hub function in the developing CA1 circuit. To this end, horizontal slices from Dlx1/2(E7,5)-GFP or wild-type mice containing the intermediate/ventral CA1 were loaded with the calcium indicator Fura2-AM and subsequently imaged using 2-photon microscopy to record spontaneous neuronal activity. As we previously reported for CA3 ebGABAs^10^, GFP+ cells could not be labeled with Fura2-AM. This prevented us from calculating their functional connectivity index based on the analysis of their spontaneous calcium events. Thus, we tested the role of ebGABAs in orchestrating spontaneous network dynamics by measuring evoked GDPs upon ebGABA stimulation.

We performed whole cell patch clamp recordings from ebGABA (GFP+ cells, *n* = 65) and ctrlGABA cells (random GABAergic cells in stratum oriens and stratum radiatum, *n* = 17, Figure 1a-b). A phasic stimulation protocol was applied, i.e., short supra-threshold current pulses repeated at 0.1, 0.2 and 0.4 Hz (within the frequency range of spontaneous GDP occurrence). Giant depolarizing potentials could be detected only in 14/82 slices (*n* = 8 ctrlGABAs, *n* = 6 ebGABAs). Thus, subsequent analyses were restricted to these cases. Stimulation of 3/6 ebGABAs significantly affected GDP frequency (among these, two decreased GDP frequency whereas one increased it, Figure 1d, g). In contrast, no stimulated ctrlGABA (0/8) significantly affected GDP frequency (Figure 1c, e and Supplementary Table 1).

**Figure 1.**
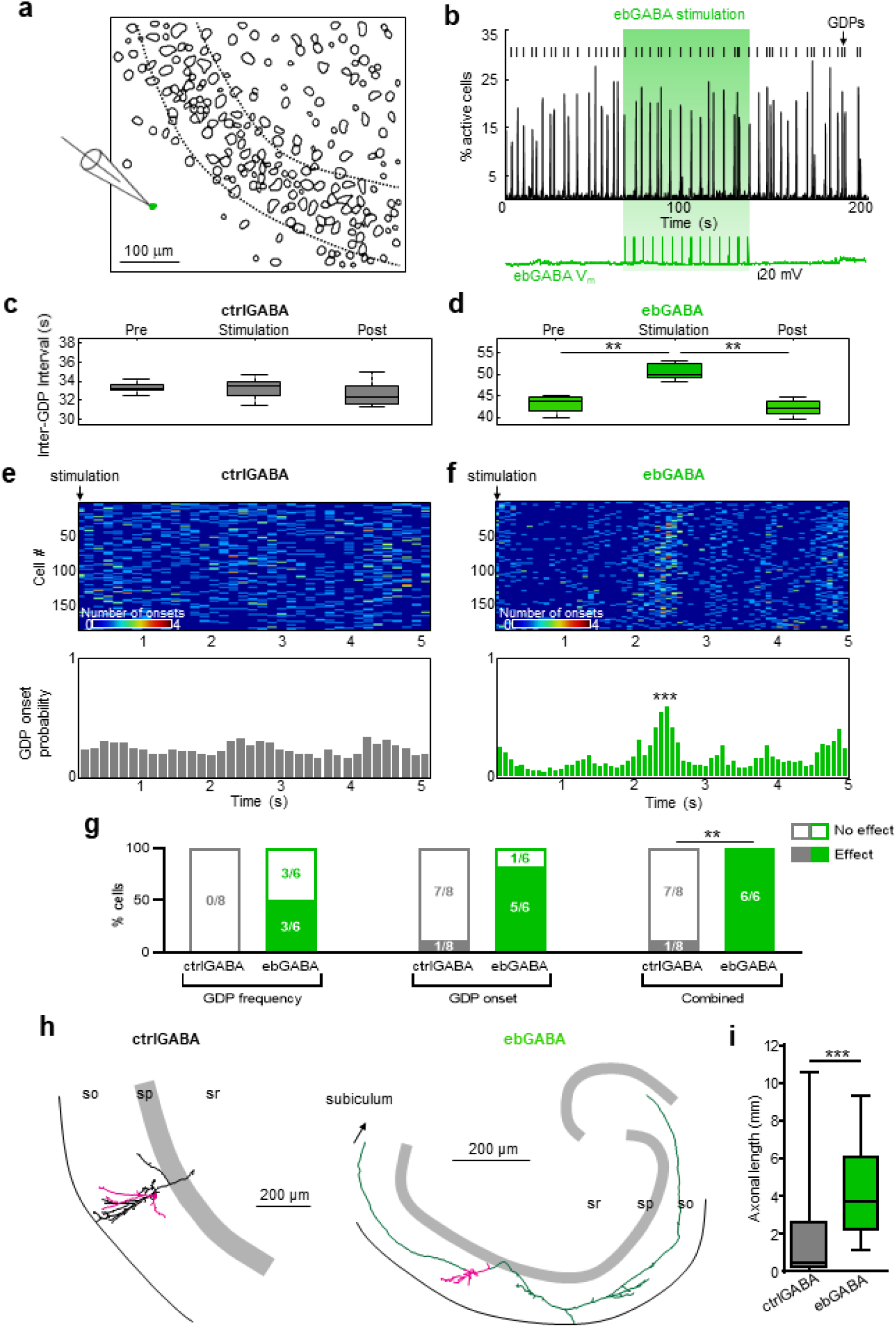
ebGABAs orchestrate network activity in the developing CA1. **(a)** Detected contours of CA1 cells imaged with 2-photon microscopy at 20 magnification. An ebGABA (green cell) localized in the stratum oriens was patched in whole cell configuration and stimulated by injecting suprathreshold depolarizing current steps. Dashed lines delimit the stratum pyramidale. **(b)** Histogram displaying the percentage of active cells in the field of view. Green area denotes stimulation of the ebGABA, the membrane potential of which is shown at the bottom. Vertical lines denote detected GDPs. **(c-d)** Box plots of ‘Inter GDP intervals’ (see Methods) of a representative ctrlGABA and a representative ebGABA cell. *(c)* stimulation of the ctrlGABA cell does not significantly affect the inter GDP interval (*P* > 0.1, Mann-Whitney U test). *(d)* stimulation of the ebGABA cell significantly increases the inter GDP interval (*P* < 0.01, Mann-Whitney U test). Data are represented as medians and interquartile ranges. Error bars represent minimum and maximum values. **(e)** Stimulation of the ctrlGABA does not synchronize GDP occurrence. *Top*, raster plot of calcium onsets following each stimulation (each row represents the average activity of single cell across 20 trials). *Bottom*, histogram showing no significantly increase in the probability of calcium events following ctrlGABA stimulation. **(f)** Stimulation of representative ebGABA synchronizes GDP occurrence. *Top*, raster plot of calcium onsets following each stimulation (each row represents the average activity of single cell across 20 trials). *Bottom*, histogram showing significantly increase in the probability of calcium events following ebGABA stimulation. **(g)** Proportions of ctrlGABA and ebGABA displaying significant effects on GDP frequency and GDP synchronization upon stimulation. A significantly higher proportion of ebGABAs impacts GDP frequency and/or synchronization (*right*, *P* = 0.0014, Fisher’s exact test). **(h)** Neurolucida reconstructions of representative neurobiotin-filled ctrlGABA and ebGABA cells. Axon is depicted in green for ebGABA and in black for ctrlGABA. Soma and dendrites are colored in magenta. **(i)** Boxplot comparing axonal lengths obtained in Neurolucida reconstructed ctrlGABA (*n* = 20) and ebGABA cells (*n* = 18). EbGABA display significantly higher axonal length than ctrlGABA (*P* = 0.0006, Mann-Whitney U test). Data are represented as medians (interquartile ranges). Boxplot whiskers represent minimum and maximum values. ** *P* < 0.01. *** *P* < 0.001

Next, we asked whether ebGABAs could also reliably synchronize the network in the shorter timescale. We examined the occurrence of calcium events in all the trials that followed the depolarizing current steps used as stimulation. We constructed a calcium event probability histogram including the activity of all the cells in the field of view and we normalized by the number of GDPs during the stimulation protocol. Within inter-stimulations trials, ebGABA activation increased synchronous calcium activity (Figure 1f). To define cells that significantly locked the onset of GDPs, we examined the first highest peak of the calcium event probability histogram during stimulation in respect to surrogate data (see Methods for details). Using this method, we calculated that 6/6 ebGABAs significantly synchronized the onset of GDPs, with GDP onset occurring 2.2 0.9 seconds after ebGABA activation (mean In contrast, only one ctrlGABA was found to significantly lock GDP onset (Figure 1e-g and Supplementary Table 1).

Overall, 1/6 ebGABA increased the frequency of GDPs and synchronized GDP occurrence, 2/6 ebGABAs decreased the frequency of GDPs and synchronized GDP occurrence, and 3/6 ebGABAs synchronized GDP occurrence but had no effect on GDP frequency. In contrast, only 1/8 ctrlGABA synchronized GDP occurrence. The remaining ctrlGABAs had no effect on either GDP frequency or onset (Supplementary Table 1). On balance, all ebGABAs could single-handedly affect the coordination of neuronal activity in the developing CA1 network, whereas the proportion of ctrlGABA affecting network dynamics (1/8) was significantly lower (*P* = 0.0014, Fisher’s exact test). Hence, CA1 ebGABAs can be classified as operational hub neurons because they display high effective connectivity, i.e. they can synchronize the activity of large population of cells in the network.

### EbGABA display widespread axonal arborization in the developing CA1

Previously described hub cells and ebGABAs in the CA3 area could be distinguished by four times longer axonal lengths compared to non-hub cells^10, 15^. We asked whether widespread axonal arborization could also be a distinctive feature of CA1 ebGABAs. To test this, we reconstructed 38 neurobiotin-filled neurons (ctrlGABAs: *n* = 20, ebGABAs: *n* = 18, Figure 1h and Supplementary Figure 1).

The axons of most CA1 ebGABAs displayed remarkable length, in some cases crossing CA1 boundaries and innervating CA3 and/or the subiculum (Figure 1h and Supplementary figure 1). In four cases, the axons ran in the alveus, suggesting a possible extrahippocampal projection. We performed morphometric analysis on the reconstructed cells. In line with previous results on CA3 ebGABA and hub cells, CA1 ebGABAs showed significantly longer axons (*P* = 0.0006) that covered a significantly larger surface than ctrlGABAs (*P* = 0.0003, Mann-Whitney U test, Figure 1i and Supplementary Figure 2a). In contrast, dendritic length or surface did not differ significantly between the two groups (*P* = 0.279 and *P* = 0.125, respectively, Mann-Whitney U test, Supplementary Figure 2b). When we pooled cells with an operational hub role (six ebGABAs and one ctrlGABA), we found that hub cells had significantly longer axons (but not dendrites) than non-hub cells (seven ctrlGABA, *P* = 0.0379, Mann-Whitney U test, Supplementary Figure 2c-d), pointing towards a link between widespread axons and a functional hub role. Thus, CA1 ebGABA exhibit functional and anatomical features of previously reported hub cells^10, 15, 16^.

### Adult ebGABAs exhibit features of long-range projecting cells

Given that ebGABAs operate as hub cells in the immature CA1 we next sought to investigate their fate in the adult CA1. We first examined their distribution across CA1 layers in PFA-fixed sections (*n* = 6 brains). The location of ebGABAs appeared to be significantly affected by layering (*P* < 0.0001, Friedman test, Figure 2a). The somata of most ebGABAs were located in the stratum oriens and inside or around the stratum pyramidale (29 ± 5% and 43 ± 4% respectively, means ± standard deviations). Fewer cells (21 ± 4%) were located in the stratum radiatum and only a small minority (7 ± 4%) populated the stratum lacunosum-moleculare. We also examined the distribution of ebGABAs in the rostrocaudal and dorsoventral axes of the hippocampus (n = 2 brains, Supplementary Figure 4). In both brains, these cells were sparse and scattered in a relatively even fashion across the entire CA1 region.

**Figure 2.**
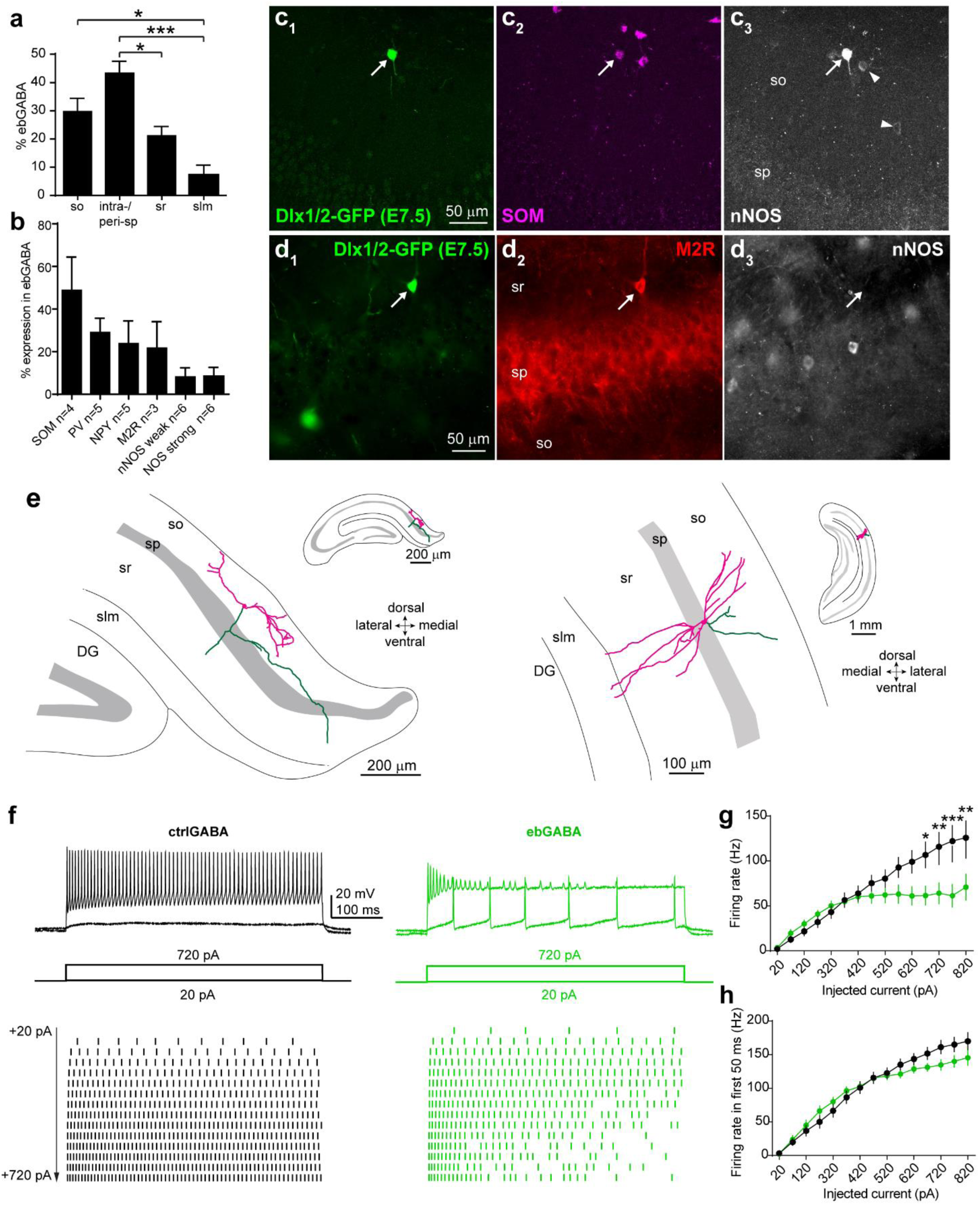
Adult ebGABAs display distinct axonal arborizations and low intrinsic firing rates. **(a)** Significant effect of layering in the proportional distribution of CA1 ebGABAs (*P* < 0.0001, *n* = 6 brains, Friedman test; so vs intra-/peri-sp *P* > 0.99, so vs sr *P* > 0.99, so vs slm *P* = 0.0439, intra-/peri-sp vs sr *P* = 0.0439, intra-/peri-sp vs slm *P* = 0.0003, sr vs slm *P* > 0.99, *post hoc* Dunn’s multiple comparisons test). **(b)** Quantification of the expression of single molecular markers in CA1 ebGABA. **(c)** Example ebGABA in stratum oriens expressing SOM and strong levels of nNOS (arrow). Note that neurons expressing weak levels of nNOS (arrowheads) are SOM immunonegative. **(d)** EbGABA in stratum radiatum expressing only M2R but not nNOS (arrow). **(e)** Neurolucida reconstructions of representative neurobiotin-filled ebGABA in the adult CA1. Axon is depicted in green. Soma and dendrites are colored in magenta. *Inset*: zoomed out representation showing the entire hippocampus to illustrate the rostro-caudal and dorso-ventral position of the neurons. **(f)** Representative responses to small and large positive current injections in ctrlGABA and ebGABA. Note the depolarization block induced in ebGABA upon strong current injection (top trace). **(g)** f/I plot showing reduced excitability of ebGABA with strong current injections. Effect of current injection F (14, 864) = 31.59, *P* < 0.0001; effect of birth date F (7, 864) = 16.40, *P* < 0.0001, interaction F (98, 864) = 1.881 *P* < 0.0001, two-way ANOVA with Bonferroni *post hoc* test. **(h)** f/I plot limited to the first 50 ms of the current injection. Effect of current injection F (14, 461) = 77.8, *P* < 0.0001; effect of birth date F (1, 461) = 3.211, *P* = 0.0738, interaction F (1, 461) = 1.676, *P* = 0.0573, two-way ANOVA with Bonferroni *post hoc* test. Data are represented as means ± standard errors of the means. * *P* < 0.05; ** *P* < 0.01; *** *P* < 0.001. so: stratum oriens; sp: stratum pyramidale; sr: stratum radiatum; slm: stratum lacunosum-moleculare.

We examined the molecular content of CA1 ebGABAs to infer the putative cell types comprising this GABAergic population. Staining for single neurochemical markers, we found that many ebGABAs expressed SOM (49 ± 16%, mean ± standard deviation) and, in a progressively lower extent, PV (29 ± 7%), NPY (24 ± 11%) and M2R (22 ± 12%, Figure 2b and Supplementary Figure 3a-b). These data are in line with previously published results on the whole hippocampus^10^. Using an antibody that allows discrimination between weak and strong levels of nNOS expression, we found that a small but consistent proportion of ebGABAs (8 ± 4%) expressed strong nNOS levels, a marker of long-range projection cells^17^ (Figure 2c and Supplementary Figure 3c).

To estimate the cell types comprised in the ebGABA population, we examined the combinatorial expression of molecular markers in these cells. In the hippocampus, strong nNOS+ cells are GABAergic projection cells that also express SOM, NPY and M2R^14^ and are likely to innervate the dentate gyrus and CA3^18^. We confirmed that strong nNOS+ ebGABAs belong to this cell type by finding that these cells often co-expressed SOM (94 ± 11%, *n*=4 brains), NPY (93 ± 13%, *n*=5 brains) and M2R (62 ± 27%, *n*=3 brains, Figure 2c and Supplementary Figure 3). EbGABAs in stratum oriens often expressed SOM and NPY but rarely PV, confirming that the majority of ebGABAs in this layer are not O-LM cells. Furthermore, some ebGABA (15 ± 7%, n=3 brains) expressed M2R but not nNOS (Figure 2d), suggesting that some of these cells could be retrohippocampal projection cells innervating the subiculum and the retrosplenial cortex^19^.

Based on this combinatorial expression of molecular markers and previous data^10, 12^, we estimated that at least 40% of ebGABA are likely to be long-range GABAergic projection cells (see Methods for details). As it was estimated that CA1 GABAergic projection cells account for only 5-7% of all GABAergic cells^20^, these data suggest that ebGABAs are biased towards a long-range output connectivity.

To corroborate these data, we filled ebGABAs with neurobiotin using whole cell patch clamp in acute coronal brain slices (*n* = 34). A subset of these cells (*n* = 14) were morphologically reconstructed to reveal their axonal and dendritic arborization (Figure 2e and Supplementary figure 5). In line with the immunohistological data, all reconstructed ebGABAs displayed little local axonal arborization, suggesting that most of these neurons do not belong to canonical interneuron types. In addition, the axons of some ebGABAs consisted of only one or two branches running straight for several hundreds of micrometers. Finally, not only the somata of ebGABAs were rarely located in stratum lacunosum-moleculare, but also ebGABAs’ dendrites arborized very little in this layer. Taken together, these data indicate that ebGABAs are biased towards a deep radial location and long-range outputs.

### Adult CA1 ebGABAs display low intrinsic firing rates

We then wished to determine whether ebGABAs mature into a physiologically defined subpopulation of GABA neurons. To this end, we performed a series of *ex vivo* whole cell patch clamp recordings in acute brain slices from adult Dlx1/2(E7.5)-GFP mice (*n* = 167 cells from 73 mice; 85 ctrlGABAs, 82 ebGABA). First, we analyzed ebGABAs intrinsic electrophysiological properties. Analysis of firing rate *versus* injected current (f/I) curves revealed a sublinear input/output relationship in ebGABA but not in ctrlGABA (Figure 2f-g; effect of current injection *P* < 0.0001; effect of birth date *P* < 0.0001, interaction *P* < 0.0001, two-way ANOVA). In many cases, upon strong current injections ebGABA displayed initial high frequency firing, but then inactivated, likely due to depolarization block^21^.

When we plotted an f/I curve only for a shorter time window of the current injection (the initial 50 ms), the firing of ebGABA was not significantly lower than the one of ctrlGABA (effect of current injection *P* < 0.0001; effect of birth date *P* = 0.0738). The remaining intrinsic membrane parameters that we examined did not differ between ctrlGABA and ebGABA, with the exception of the sag ratio, which was higher in ebGABA (Supplementary Table 2). Thus, these data suggest that ebGABA display lower firing rates than ctrlGABA only for long and strong depolarizing inputs, but not for short ones.

### Synaptic excitation of ebGABAs is biased towards intra-hippocampal connections

Since the somata of ebGABAs are preferentially localized in deep CA1 layers and CA1 inputs are radially organized, we asked whether an early birth date biases GABAergic cells towards input connectivity from glutamatergic pathways. We probed ebGABAs’ monosynaptic input connectivity from specific pathways using electrical stimulation and optogenetic mapping combined with intracellular voltage clamp recordings (with cells held at ECl: -87 mV). First, we focused on the Schaffer collateral input by stimulating the axons from CA3 pyramidal cells in stratum radiatum using a bipolar stimulating electrode while recording from CA1 GABA cells in voltage clamp (Figure 3a). Electrical stimulation of the Schaffer collateral evoked EPSCs in the majority of ctrlGABAs (13/16 cells) and ebGABAs (9/14 cells, Figure 3b-c). EPSCs were mediated by AMPA/KA receptors because they were blocked by NBQX (10 µM, 3/3 cells, Supplementary Figure 6). Thus, the majority of ebGABAs are recruited by CA3 inputs.

**Figure 3.**
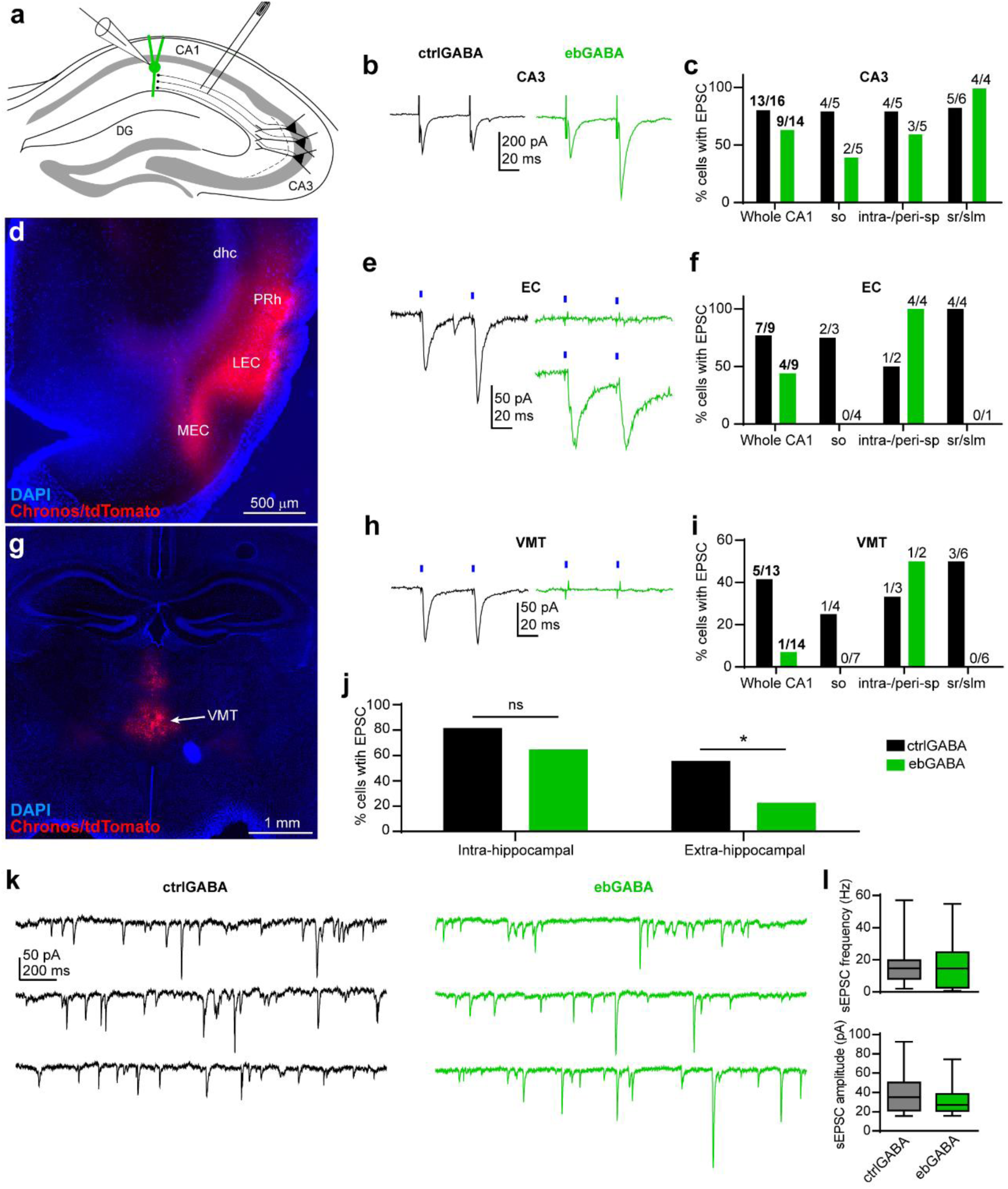
Sparse extra-hippocampal excitatory input connectivity onto ebGABAs. **(a)** Schematic of the experimental configuration: a bipolar tungsten electrode was positioned in the stratum radiatum between CA1 and CA3 to stimulate the Schaffer collateral pathway. **(b)** Representative EPSCs evoked by electrical stimulation of the Schaffer collateral in a ctrlGABA and an ebGABA cell. Stimulation artifacts were trimmed for clarity. **(c)** Proportions of CA1 ctrlGABA and ebGABA cells in which EPSCs could be evoked upon electrical stimulation of the Schaffer collateral. **(d)** representative injection site for EC stimulation experiments. Chronos/tdTomato expression is mostly restricted to the lateral entorhinal cortex (LEC). MEC: medial entorhinal cortex. PRh: perirhinal cortex. dhc: dorsal hippocampal commissure. **(e)** Representative EPSCs evoked by optical stimulation of EC terminals. Representative ctrlGABA cell responding to the stimulation (*left*), and two representative ebGABAs: one showing no detectable response (*top right*) and one showing light-evoked EPSCs (*bottom right*). **(f)** Proportions of CA1 ctrlGABA and ebGABA cells in which EPSCs could be evoked upon optical stimulation of EC axons. **(g)** representative injection site for VMT stimulation experiments. Chronos/tdTomato expression is mostly restricted to nuclei rhomboid and reuniens of the VMT. **(h)** Representative EPSCs evoked by optical stimulation of VMT terminals. Representative ctrlGABA cell responding to the stimulation (*left*), and a representative ebGABA cell displaying no detectable response (*right*). **(i)** Proportions of CA1 ctrlGABA and ebGABA cells in which EPSCs could be evoked upon optical stimulation of VMT axons. **(j)** The proportion of ctrlGABA and ebGABA receiving intra-hippocampal excitatory inputs (CA3) was not significantly different (*P* = 0.417, Fisher’s exact test, 13/16 ctrlGABA, 9/14 ebGABA). In contrast, the proportion of ebGABA receiving extra-hippocampal excitatory inputs (pooled EC and VMT data) was significantly lower than the proportion of ctrlGABA (*P* = 0.0331, Fisher’s exact test, 12/22 ctrlGABA, 5/23 ebGABA) **(k)** Example voltage clamp traces recorded from a ctrlGABA cell and an ebGABA cell at -87 mV (E_Cl-_) to isolate sEPSCs. **(l)** Boxplots showing similar sEPSC frequency (*P* = 0.4, Mann-Whitney U test) and amplitude (*P* = 0.42, Mann-Whitney U test) in (n = 18) and ebGABA (*n* = 17). Data are represented as medians (interquartile range). Boxplot whiskers represent minimum and maximum values. * *P* < 0.05.

Since somata and dendrites of ebGABAs are rarely found in the stratum lacunosum-moleculare, we predicted that long-range inputs targeting this layer would play a less important role in the recruitment of these cells. To evaluate this hypothesis, we tested the inputs from the entorhinal cortex (EC) and the ventromedial thalamus (VMT) using a viral vector to express the fast opsin Chronos in these regions (Figure 3d-i). Following at least two weeks of expression, Chronos/tdTomato+ axons densely innervated the stratum lacunosum-moleculare of CA1 for both injection sites (some axons were also detected in the stratum oriens next to the alveus, Supplementary Figure 6; very few axons were also present in other layers). Pulses of 470 nm light were delivered to test for the presence of short-delay EPSCs (monosynaptic connections) in the recorded cells. For EC stimulations, we detected EPSCs in the majority of ctrlGABAs (7/9 cells), but only in about half of the ebGABAs (4/9 cells, Figure 3e-f). For VMT stimulations, we detected EPSCs in 5/13 ctrlGABAs cells but only in 1/14 ebGABAs (Figure 3h-i). The proportion of ebGABA recruited by an intra-hippocampal excitatory input (CA3) was not significantly different from the proportion of ctrlGABA (*P* = 0.417, Fisher’s exact test). In contrast, the proportion of ebGABA recruited by extra-hippocampal excitatory inputs (EC and VMT pooled) was significantly lower than the proportion of ctrlGABA (5/23 and 12/22, respectively, *P* = 0.0331, Fisher’s exact test, Figure 3j). When we compared the maximum amplitude and paired pulse ratio for CA3, EC and VMT inputs, we observed similar values in the two cell populations (Supplementary Figure 6f-g), indicating that an early birth date biases GABA cells’ input connectivity but not synaptic strength or release probability. Given the sparse connectivity and the fact that EPSCs evoked in ebGABAs by stimulation of the EC were small (maximum amplitude 19 ± 33 pA, mean ± standard deviation), these data suggest that extra-hippocampal afferents play a small role in the recruitment of ebGABAs.

Furthermore, we examined spontaneous EPSCs (sEPSCs), many of which are likely to be action potential-dependent, and thereby arise from intra-hippocampal connections in coronal slices. We confirmed that sEPSCs were mediated by AMPA/KA receptors because they were completely blocked by NBQX (10-20 µM, 4/4 cells). CtrlGABA and ebGABA displayed similar sEPSC parameters, in particular frequency and amplitude (Figure 3k-l and Supplementary Figure 6h), indicating similar recruitment of these cells by putative intra-hippocampal excitatory afferents. Taken together, these data indicate that the deep location and dendritic arborization of CA1 ebGABA favor their recruitment by intra-hippocampal inputs.

### ebGABAs receive weak local synaptic inhibition

Given that ebGABAs are biased towards a deep location and that the radial position influences the inhibitory innervation of CA1 pyramidal cells^22^, we also asked whether an early birth date biases GABA cells towards specific inhibitory wiring schemes. We began by examining sIPSCs by voltage clamping the cells at the reversal potential for glutamatergic currents (0 mV; ctrlGABA *n* = 18, ebGABA *n* = 17, Figure 4a). As many sIPSCs are action potential-dependent, this measurement largely portrays local inhibition driven by interneurons firing in the slice. We confirmed that sIPSCs were mediated by activation of GABAA receptors as they were completely abolished by the GABAA receptor antagonist Gabazine (SR95531, 10 µM, *n* = 4 cells). EbGABA displayed a significantly lower sIPSC frequency compared to ctrlGABA (ctrlGABA: median 11.7 Hz, IQR 6.2 – 16.8 Hz; ebGABA: median 6.5 Hz, IQR 1.2 – 10.6 Hz; *P* = 0.043, Mann Whitney U test). In contrast, other sIPSC parameters were comparable between ctrlGABA and ebGABA (Supplementary Figure 7a). Lower frequency of sIPSCs in ebGABA could arise from decreased spontaneous activity of interneurons innervating ebGABA, sparser innervation by GABAergic terminals or lower release probabilities of presynaptic GABAergic terminals. To verify whether weaker innervation by GABAergic terminals underlies this effect, we carried out further experiments.

**Figure 4.**
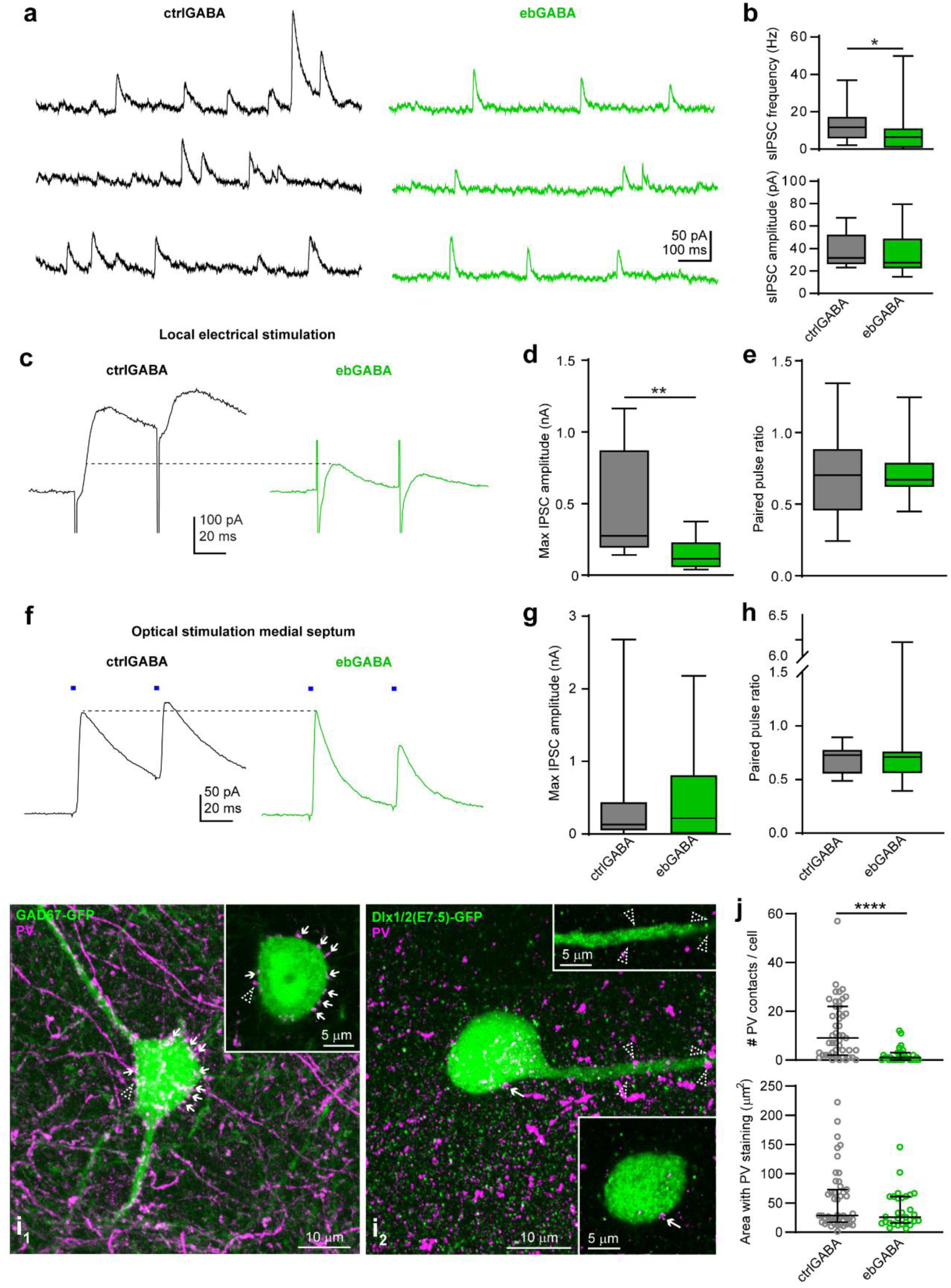
Sparse local inhibitory inputs onto adult ebGABAs. **(a)** Example voltage clamp traces recorded from a ctrlGABA and an ebGABA at 0 mV (E_EPSC_) to isolate sIPSCs. sIPSC frequency is lower in the ebGABA cell. **(b)** sIPSC frequency but not amplitude is significantly lower in ebGABAs (*n* = 17) than ctrlGABAs (*n* = 18; *P* = 0.043, Mann Whitney U test). **(c)** Representative IPSCs evoked by local electrical stimulation (stimulating electrode placed in the stratum radiatum). IPSCs evoked in the ctrlGABA cell display higher amplitude than the ones evoked in the ebGABA cell. Stimulation artifacts were trimmed for clarity. **(d)** The maximum amplitude of the IPSC evoked by local electrical stimulation is significantly lower in ebGABAs than ctrlGABAs (*P* = 0.0085, Mann-Whitney U test, ctrlGABA *n* = 10, ebGABA n = 8). **(e)** IPSC paired pulse ratio is not statistically different between ctrlGABAs and ebGABAs (*P* = 0.792, Mann-Whitney U test). (**f)** representative IPSCs evoked by optical stimulation of medial septal terminals. IPSCs evoked in ctrlGABA and ebGABA cells display similar amplitudes. **(g-h)** No significant difference between ebGABAs (*n* = 12) and ctrlGABAs (*n* = 13) in maximum amplitude (*P* = 0.611) and paired pulse ratio (*P* = 0.796, Mann-Whitney U tests) of the IPSCs evoked by optical stimulation of septal terminals. Data are represented as medians (intequartile ranges). **(i)** Representative maximum intensity projections of confocal z-stacks showing innervation of a ctrlGABA and an ebGABA by PV+ boutons. Appositions between PV+ boutons and GFP+ somata or dendrites are highlighted with arrows; lack of apposition is marked with dashed arrowheads . *Insets*: single optical sections (thickness 0.41 µm) showing magnifications of the appositions. (*i_1_*): z-stack depth 20.5 µm. (*i_2_*): z-stack depth 15.17 µm. **(j)** EbGABAs display a significantly lower number of PV+ appositions than ctrlGABAs (*top*, *P* < 0.0001, Mann-Whitney U test); area of the PV staining is not significantly different between the two populations of cells (*bottom*, *P* = 0.236, Mann-Whitney U test, ctrlGABA *n* = 47, ebGABA *n* = 30). Horizontal lines represent quartiles. * *P* < 0.05. ** *P* < 0.01. **** *P* < 0.0001.

First, we placed a bipolar stimulating electrode in stratum radiatum, the layer with weakest innervation by medial septal axons (Supplementary Figure 7c) to bias our stimulation towards interneurons. In line with the sIPSC data, the maximum IPSC evoked by local electrical stimulation was significantly higher in ctrlGABA than ebGABA (*P* = 0.0085, Mann-Whitney U test, ctrlGABA n=10, ebGABA n = 8, Figure 4c-d). In contrast, IPSC paired pulse ratio did not differ between the groups (Figure 4e). These observations suggest that sparser GABAergic innervation and not weaker release probability at GABAergic synapses could account for the lower inhibitory tone onto ebGABA.

To corroborate that this deficit in inhibition arises from local sources, we probed the GABAergic input from the medial septum, a region that provides significant innervation of CA1 GABAergic cells^23^, by viral Chronos expression in this nucleus (Supplementary Figure 6a-b). Following at least two weeks of expression, Chronos/tdTomato+ axons densely innervated the stratum oriens and the border between the stratum radiatum and the stratum lacunosum-moleculare (and to a lesser extent the strata pyramidale and radiatum; Supplementary Figure 7c). Both ctrlGABA and ebGABA displayed a high degree of connectivity (13/14 ctrlGABA and 12/18 ebGABA receiving IPSCs upon septal stimulation, Supplementary Figure 6e). These IPSCs were GABAergic as they were blocked by the GABAA receptor antagonist Gabazine (SR95531, 10 µM, 11/11 cells, Supplementary Figure 7d). Maximum amplitude of the IPSC evoked by septal stimulation as well as paired pulse ratio did not differ between ctrlGABA (n = 13) and ebGABA (n = 12, Figure 4f-h), suggesting that these cells received similar innervation and release probability from this pathway. Thus, the deficit in inhibition onto ebGABAs is likely to originate from local sources.

### Sparse innervation by parvalbumin-positive cells underlies weak inhibition in ebGABAs

Since parvalbumin-expressing (PV+) basket cells were previously shown to differentially target CA1 pyramidal neurons according to their radial position (Lee et al 2014), we tested whether reduced innervation by PV+ cells could account for the low local inhibition observed. To this end, we quantified the number of PV+ boutons innervating ebGABAs and ctrlGABAs. To quantify the innervation ctrlGABAs, we used the GAD67-GFP mouse line. To avoid bias due to uneven sampling across layers, we sampled ctrlGABA cells that roughly matched the position of the imaged ebGABAs (*n* = 43 ctrlGABA and *n* = 30 ebGABA, each population from 2 brains). We found a striking difference in the number of boutons innervating the two populations, with ebGABAs receiving a significantly lower number of PV+ terminals (*P* < 0.0001, Mann Whitney U test, Figure 4i-j). Importantly, we verified that intensities and areas of the PV stainings, as well as the sampled volumes were similar for the two populations (*P* = 0.2363, Mann-Whitney U test, Figure 4j and Supplementary Figure 8). Additionally, the difference in the number of PV+ terminals held when restricting the analysis to cells in the stratum oriens (*P* = 0.0008, Mann-Whitney U test, Supplementary Figure 8d), suggesting that this difference was not generated by uneven sampling across layers. Thus, an early birth date biases GABA cells for a sparse PV innervation, which is likely to account (at least in part) for the weak inhibition observed in ebGABAs.

### ebGABAs are highly recruited by locomotion and network events *in vivo*

Finally, we asked whether ebGABAs’ different anatomical, electrophysiological and wiring properties reported above could result in distinct *in vivo* activity in awake mice. To test this, we injected a viral vector expressing the red calcium indicator jRGECO1a in the dorsal hippocampus of adult Dlx1/2(E7.5)-GFP mice. Two weeks after the injections, mice were implanted with a chronic glass window that was placed just above the dorsal hippocampus and a bar for head fixation. This allowed performing *in vivo* 2-photon calcium imaging from ebGABA and nearby cells. Mice were head-fixed in the dark on a non-motorized treadmill allowing spontaneous movement (Figure 5a, Villette et al., 2015a).

We imaged from 39 mice expressing jRGECO1a in large numbers of cells using 400 × 400 µm fields of view. Given the sparseness of ebGABAs, the field of view contained an ebGABA only in 3/39 mice (one cell in the stratum oriens, two cells in the stratum pyramidale, Figure 5b). The calcium dynamics of these three ebGABAs and nearby cells (24-102 cells) were imaged during spontaneous locomotion and rest. First, we examined single cells’ responses to locomotion (Figure 5d, see Methods for details). We found that all three ebGABAs were part of a minor-to-moderate proportion of cells that were consistently recruited by locomotion (Locomotion_ON_ cells, 12-52%, Figure 5f, j). Second, we analyzed single cell activities in relation to synchronous calcium events (SCEs, Figure 5e, see Methods for details) that are known to occur during rest, often in synchrony with sharp-wave ripples^25^. SCEs occurred at a rate of 0.09-0.16 Hz in stratum pyramidale, in line with previous reports, and of 0.03 Hz in stratum oriens. We found that all three ebGABAs were part of a majority of CA1 cells that were significantly recruited at least once during SCEs (SCE_ON_ cells, 67-86%, Figure 5g, j). Third, we detected cell assembly patterns occurring during rest in a 200 ms time window and analyzed single cell activities in relation to assembly activations (Figure 5h, see Methods for details). This analysis revealed that ebGABAs were part of a minority of cells that were significantly recruited by cell assemblies (Assembly_ON_ cells, 15-37%, Figure 5i-j). Overall, ebGABAs were among a minority of imaged cells that were significantly recruited by locomotion, SCEs and assembly activities (highly recruited cells, 3-26%, Figure 5j). This suggests that ebGABAs are highly involved in a broad range of hippocampal dynamics during both locomotion and rest.

**Figure 5.**
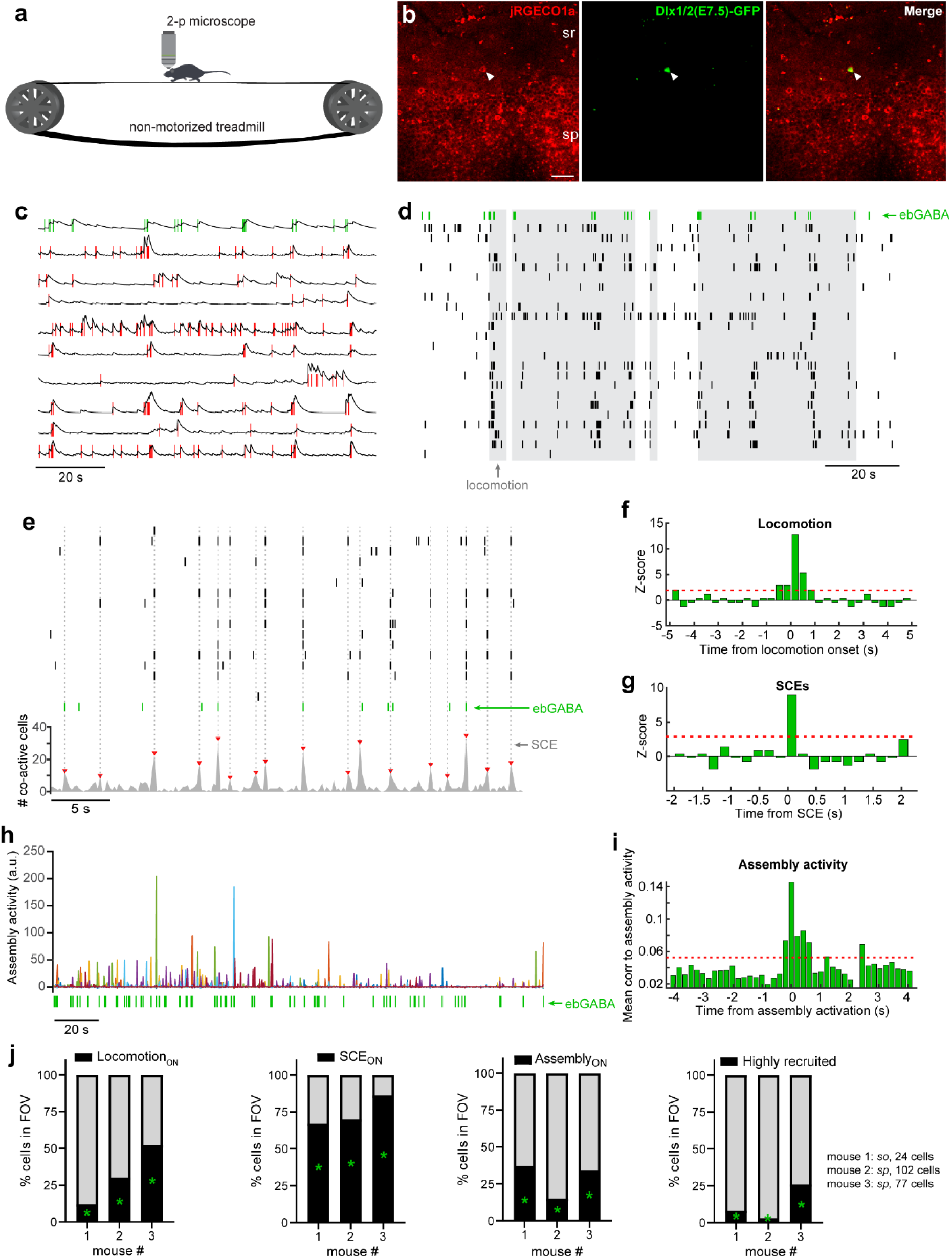
Adult ebGABAs are recruited by locomotion and network activity *in vivo*. **(a)** Schematic of the experimental configuration. **(b)** Two-photon *in vivo* jRGECO1a and Dlx1/2(E7.5)-GFP signals imaged from the stratum pyramidale of CA1 (mouse 2). Scale bar: 50 µm. **(c)** 20% ΔF/F traces extracted from somata in the field of view shown in b and corresponding spikes (vertical lines) estimated with the CaImAn toolbox. The top trace with green spikes was extracted from the ebGABA (GFP-positive) cell, whereas the remaining traces were extracted from the somatic signal of GFP-negative cells. **(d)** Raster plot showing the activity of the ebGABA and 23 GFP-negative cells in response to locomotion (gray areas). **(e)** *Top*, raster plot showing the activity of the ebGABA and 19 GFP-negative cells in response to SCEs (gray vertical lines). *Bottom*, number of co-active cells in time. Red dashed lines denote a significant SCE. **(f)** Z-score showing a significant response to locomotion of the ebGABA. **(g)** Z-score showing a significant response to SCEs of the ebGABA. **(h)** Expression of assembly activities in time during rest (see Methods for details). Seven significant assemblies were detected (assemblies are color coded). **(i)** Mean cross-correlogram between ebGABA spike count and the activity of each assembly showing a correlation above chance level (see Methods for details). Panels c-i refer to the same field of view shown in b. **(j)** Proportion of cells recruited by locomotion (Locomotion_ON_ cells), SCEs (SCE_ON_ cells), assembly activity (Assembly_ON_ and all three events combined (highly recruited cells). Green asterisks denote the group to which the ebGABA cell belongs.

## Discussion

Using a combination of techniques including inducible genetic fate mapping, *ex vivo* and *in vivo* large-scale calcium imaging, electrophysiology, optogenetics, and anatomical analyses, we have shown that ebGABAs are involved in local CA1 dynamics both in development and adulthood. At the neonatal stage, ebGABAs coordinate network bursts (GDPs) *ex vivo*. In adulthood, ebGABAs signal a variety of network states *in vivo*. Their early birth date specifies anatomical and intrinsic firing properties as well as input connectivity schemes that likely to contribute to their recruitment.

Remarkably, ebGABAs maintain a set of distinct anatomical and functional properties in the adult CA1. EbGABAs display sparse local axons and express combinations of markers typical of different classes of projection cells. Electrophysiologically, they are characterized by a sublinear f/I (input/output) relationship limiting strong activations to short periods of time. Additionally, adult ebGABAs show wiring schemes that are distinct from randomly sampled GABAergic cells. Their recruitment appears to be largely driven by intra-hippocampal excitatory afferents because they receive typical intra-hippocampal excitation, but little long-range inputs from the entorhinal cortex and the ventromedial thalamus. Furthermore, ebGABAs receive weak local inhibition via a sparse innervation by axons arising from PV+ neurons. *In vivo*, ebGABAs are part of a minority of cells in the CA1 network that are recruited by locomotion, synchronous calcium events and assembly activity. Therefore, ebGABAs appear predetermined for exceptional functional and structural properties in both the developing and adult hippocampus.

The present study demonstrates that ebGABAs are highly involved in hippocampal network dynamics in development and adulthood. During early postnatal development, stimulation of a single ebGABA is sufficient to change the frequency of GDPs and to reliably trigger their onsets. The latter phenomenon is likely to involve complex polysynaptic interactions because the delay between ebGABA stimulation and GDP onset ranged between 0.2 and 5 seconds. Thus, hub cells are not a unique feature of CA3, a highly recurrent network^26^, but are distributed throughout the hippocampal formation, and possibly throughout the brain. It is not clear whether CA1 hub cells orchestrate GDPs only through action on the CA1 circuit or, by contrast, they modulate GDP generation in CA3 with subsequent propagation to CA1. Both scenarios are possible. Morphological reconstructions of biocytin-filled ebGABA demonstrated that some projected back to CA3, a finding that is corroborated by our analyses of molecular marker combinations (i.e. strong nNOS-expressing backprojection cells). On the other hand, GDPs have been shown to occur in CA1 also independently from CA3^13, 27^.

Giant depolarizing potentials and their *in vivo* counterpart^28^ are network bursts that are thought to represent an early form of sharp wave ripple (SWRs). Sharp wave ripples are generated in CA2 and, similarly to GDPs, require the recurrent circuit of CA3 to successfully propagate to CA1^29, 30^. Recently, we reported that SCEs detected with calcium imaging in the adult CA1 often occur during SWRs and involve reactivations of cell assemblies^25^. Here we show that ebGABAs maintain a strong relationship with network bursts (SCEs) in adulthood. Additionally, these cells are consistently recruited around the activation of CA1 assemblies. This may be achieved via special intrinsic properties and circuit motifs encompassing ebGABAs. For instance, we have shown with *ex vivo* patch clamp experiments that ebGABAs display efficient rate coding for short but not long depolarizing stimuli, a mechanism that could favor their transient recruitment *in vivo*. In addition, our data suggest that ebGABAs receive the majority of their excitatory inputs from intra -hippocampal connections. This may enhance their signal to noise ratio to report local network activity to postsynaptic targets. Finally, we demonstrate that ebGABAs receive little local inhibition from PV+ interneurons, but high levels of long-range inhibition from the medial septum. This finding is consistent with a study showing that the majority of inhibitory terminals on CA1 GABAergic projection cells arise from the medial septum^31^. Virtually all PV+ terminals originating from CA1 interneurons and targeting other GABA cells are likely to arise from PV+ basket cells and bistratified cells, and both cell types are strongly active during SWRs^32, 33^. Thus, this lack of cell type-specific inhibition could additionally favor ebGABAs’ recruitment during SWRs/SCEs.

Another remarkable feature of ebGABAs in the adult CA1 *in vivo* is their functional versatility, namely their recruitment during a variety of behavioral/network phenomena. Specifically, ebGABAs were among a minority of cells in the network that were consistently activated during locomotion, SCEs and assembly activations. This versatility sets them apart from most known GABAergic cell types because interneurons that are activated by locomotion are usually not activated during SWRs, and vice versa^32–34^.

Interestingly, ebGABAs’ recruitment during a variety of network states is reminiscent of the activity described for CA1 GABAergic projection neurons^35^. In line with this, a significant portion of ebGABAs project to the medial septum^12^. In the present study, we have found that ebGABAs express combinations of molecular markers typical of two extra classes of GABAergic projection cells: strong nNOS-expressing backprojection cells, likely innervating dentate gyrus and CA3^14, 18^, and M2R-expressing retrohippocampal cells innervating subiculum and retrosplenial cortex^19^. Combined with the fact that biocytin-filled ebGABAs exhibited little local axonal arborization, these data indicate that ebGABAs could form several classes of projection cells innervating various target regions.

We report that an early embryonic origin results in a deep soma location, sublinear f/I relationship and reduced innervation by PV+ interneurons and by long-range excitatory inputs, in particular by thalamic afferents. The putative depolarization block observed in ebGABA could result from the interplay between sodium and delayed rectifier potassium conductances^21^. Interestingly, a similarly lower intrinsic excitability was recently reported to distinguish granule cells with an early temporal embryonic origin^36^, suggesting that this may represent a common feature of early born cells, regardless of the neurotransmitter that they release.

An early embryonic origin specifies a large proportion of long-range projecting GABAergic cells, an extremely rare neuronal population^20^. It is crucial to understand why and by which mechanisms this early specification occurs. It was recently proposed that the transcription factor CoupTF2 may act as a temporal identity cue that promotes an early specification towards somatostatin-expressing cortical GABA neurons^37^, suggesting that the majority of ebGABAs may well also critically depend on this transcription factor. Future studies are needed to further determine the identity and spatial location of ebGABAs progenitor cells within the embryonic telencephalon. We hypothesize that early transcription factors shared across different subpallial proliferative areas could direct GABAergic cells towards a long-range projecting fate.

On balance, this study shows that ebGABAs are pioneer GABAergic cells operating as “hubs” during development and maintaining unique features throughout adulthood. To our knowledge, we have provided the first evidence that an early birth date alone (regardless of spatial embryonic origins or cell types) dictates anatomical, electrophysiological and connectivity properties of GABAergic cells. Given their bias towards long-range targets, intra-hippocampal inputs and local assembly activity, we hypothesize that ebGABAs could detect CA1 activity and bind local and distant assemblies into chains of neuronal activation^38^.

## Methods

### Animals

All protocols were performed under the guidelines of the French National Ethics Committee for Sciences and Health report on ‘‘Ethical Principles for Animal Experimentation’’ in agreement with the European Community Directive 86/609/EEC under agreement #01413. Dlx1/2Cre^ER+/-^;RCE:LoxP^+/+^ male mice^39^ were crossed with 7-8 weeks old wild-type Swiss females (C.E Janvier, France) for offspring production. To induce Cre^ER^ activity, we administered a tamoxifen solution (Sigma) by gavaging (force-feeding) pregnant mice with a silicon-protected needle (Fine Science Tools). We used 2 mg of tamoxifen solution per 30 g of body weight prepared at 10 mg/mL in corn oil (Sigma). Pregnant females crossed with Dlx1/2Cre^ER+/-^;RCE:LoxP^+/+^ males were force-fed at embryonic days 7.5 days post vaginal plug (corresponding to the embryonic ages E7.5) in order to label early expressing Dlx1/2 precursors with GFP in the embryos. For simplicity, we refer to Dlx1/2Cre^ER+/-^;RCE:LoxP^+/+^ as Dlx1/2(E7.5)-GFP. GAD67-GFP mice were kindly donated by Dr. Hannah Monyer (Heidelberg University).

### Slice Preparation for *ex vivo* experiments

Slices containing the hippocampus were prepared from Swiss wild-type or Dlx1/2(E7.5)-GFP mice. Four to five days old (P4–P5) pups were used for developmental experiments and 4-33 weeks old mice (mean age ± SD: P67 ± 32) for adult experiments. Adult mice were anaesthetized with a ketamine/xylazine mix (Imalgene 100 mg/kg, Rompun 10 mg/kg) or a Domitor/Zoletil mix (0.6 mg/kg and 40 mg/kg, respectively) prior to decapitation. Slices were cut using a Leica VT1200 S vibratome in ice-cold oxygenated modified artificial cerebrospinal fluid with the following composition (in mM): 2.5 KCl, 1.25 NaH_2_PO_4_, 7 MgCl_2_, 5 CaCl_2_, 26 NaHCO_3_, 5 D-glucose, 126 CholineCl. Slices were cut in the horizontal plane at 380 µm thickness for developmental experiments and in the coronal plane at 300 µm thickness for adult experiments. Slices were then incubated in oxygenated normal ACSF containing (in mM): 126 NaCl, 3.5 KCl, 1.2 NaH_2_PO_4_, 26 NaHCO_3_, 1.3 MgCl_2_, 2.0 CaCl_2_, and 10 D-glucose (1 h at room temperature, RT for developmental experiments, 15 min at 33°C followed by 30 min at RT for adult experiments).

### *Ex vivo* calcium imaging and patch clamp recordings during development

For Fura2-AM loading, slices were incubated in a small vial containing 2.5 ml of oxygenated ACSF with 25 ml of a 1 mM Fura2-AM solution (in 100% DMSO) for 30 min. Slices were incubated in the dark, and the incubation solution was maintained at 30–33°C. Slices were transferred to a submerged recording chamber and continuously perfused with oxygenated ACSF (3 mL/min) at 30°C–33°C. Imaging was performed with a multibeam multiphoton pulsed laser scanning system (LaVision Biotech) coupled to a microscope as previously described ^40^. Images were acquired through a CCD camera, which typically resulted in a time resolution of 80 – 138 ms per frame. Slices were imaged using a 20×, NA 0.95 objective (Olympus). Imaging depth was on average 80 µm below the surface (range: 50–100 m).

Patch recording electrodes (4-8 MΩ resistance) were pulled using a PC-10 puller (Narishige) from borosilicate glass capillaries (GC150F-10, Harvard Apparatus) and filled with a filtered solution consisting of the following compounds (in mM): 130 K-methylSO4, 5 KCl, 5 NaCl, 10 HEPES, 2.5 Mg-ATP, 0.3 GTP, and 0.5% neurobiotin (265–275 mOsm, pH 7.3). Electrophysiological signals were amplified (EPC10 amplifier; HEKA Electronik), low-pass filtered at 2.9 kHz, digitized at 10 kHz and acquired using a Digidata 1550 Digitizer and pClamp 10 software (Molecular Devices). For most stimulation experiments, imaging acquisition was separated between: (1) a baseline period during which the cell was held close to resting membrane potential (i.e., zero current injection); (2) a stimulation period during which phasic stimulation protocols were applied; and (3) a 3 min recovery period during which the cell was brought back to resting membrane potential. The stimulation protocol consisted of suprathreshold current pulses (amplitude: 100–150 pA) of 200 ms duration repeated at 0.1, 0.2 and 0.4 Hz. Cells were discarded if they did not meet the following criteria: (1) stable resting membrane potential; (2) stable network dynamics measured with calcium imaging (i.e., the coefficient of variation of the inter-GDP interval did not exceed 1); (3) cells displaying healthy shape and good Fura2-AM loading throughout the entire field of view.

### Analysis of *ex vivo* calcium imaging data during development

We used custom designed MATLAB software allowing automatic identification of loaded cells, measurement of the average fluorescence transients from each cell as a function of time and detection of onsets and offsets of calcium signals^15^. Network synchronizations (GDPs) were detected as synchronous onsets peaks including more neurons than expected by chance, as previously described^15^, and their time stamp denoted by *t_G_*. The ‘Inter GDP intervals’ (IGI) was defined as the interval between two consecutive GDPs. To establish whether the stimulation of a single neuron was able to influence the frequency of GDPs occurrence, we employed a previously published analysis^15^. We first calculated the average IGI in the three epochs: pre-stimulus (control), during the stimulation period and post-stimulus. Due to the variability distribution of IGI in each interval, we calculated the average IGI in a window of *t_s_* (300 or 400 frames calculated starting from each *t_G_* and eliminating the data corresponding to overlaps between epochs). To test for the significance of the change in the period of GDP, a Kolmogorov-Smirnov test was applied between all the three resulting distributions of average IGI and a significance level of *P* < 0.05 was chosen. To test for changes in phase of the GDP upon stimulation, the IGI during resting conditions was used as a reference period of a harmonic oscillator mimicking the rhythm of GDPs. The phase of the expected GDP generated by the harmonic oscillator was compared to the phase of the real GDP intervals occurring during the entire recording. For the i-th GDP, a phase measure *ϕ_i_* in respect to the control IGI is defined as follows:

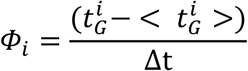

Where Δt is the average IGI interval in the control condition, and 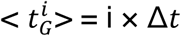 is the expected occurrence of the i-th GDP, according to the control condition. The phase of each real GDP was set to zero at the peak of synchronous activation and increased linearly reaching 2π at the peak of the next GDP. By subtracting the phase of real GDPs from the expected GDP, we could instantaneously measure the effect of phasic stimulation. The cell was included in the dataset only if the difference between expected and observed cycles did not exceed ± 2 cycles during resting conditions.

To assess whether single cell stimulation was able to lock GDP occurrence, we isolated the trials following each stimulation (trial length was either 2.5 s or 5 s). Next, we calculated an average histogram of single cell calcium onsets including all trials. The histogram was then normalized by the number of GDPs during the stimulation protocol. The height of the highest peak in the histogram was stored. To estimate the significance of the time locking, we created surrogate data consisting of concatenated control and post-stimulation frames (without stimulation trials). The same histogram was then produced from the surrogate data. Since no stimulation was present in this case, we used an arbitrary frame as stimulation time. For each cell, number of trials and trial length for the surrogate data matched the ones of stimulation data. This process was repeated 100 times using 100 different starting frames for the stimulation, and the values of the highest peaks were stored. A final histogram of peak heights was computed, and the cell was considered to significantly lock GDP onset if the peak height during stimulation fell within the 5% highest peaks in the distribution (i.e. *P* < 0.05).

### Virally-driven opsin/jRGECO1a expression

Dlx1/2(E7.5)-GFP mice (P25-P35 age for opsin expression, 2-3 months age for jRGECO1A expression) were anaesthetized using either a ketamine/xylazine mix (Imalgene 100 mg/kg, Rompun 10 mg/kg) or 1-3% isoflurane in oxygen. Analgesia was also provided with buprenorphine (Buprecare, 0.1 mg/kg). Lidocaine cream was applied before the incision for additional analgesia. Mice were fixed to a stereotaxic frame with a digital display console (Kopf, Model 940). Under aseptic conditions, an incision was made in the scalp, the skull was exposed, and a small craniotomy was drilled over the target brain region. A recombinant viral vector was delivered using a glass pipette pulled from borosilicate glass (3.5” 3-000-203-G/X, Drummond Scientific) and connected to a Nanoject III system (Drummond Scientific). The tip of the pipette was broken to achieve an opening with an internal diameter of 25-35 μm. The following viral vectors were used (both from Penn Vector Core): AAV8.Syn.Chronos-tdTomato.WPRE.bGH to drive Chronos expression in the ventromedial thalamus or medial septum, AAV1.Syn.NES.jRGECO1a.WPRE.SV40 to drive jRGECO1a expression in the dorsal CA1 (for the latter, virus stock was diluted 1:4 in D-PBS, Sigma Aldrich). Stereotaxic coordinates were based on a mouse brain atlas (Paxinos and Franklin). All coordinates are in millimeters. Anteroposterior (AP) coordinates are relative to bregma; mediolateral (ML) coordinates are relative to the sagittal suture; dorso-ventral (DV) coordinates are measured from the brain surface. For entorhinal cortex, ventromedial thalamus (nuclei reuniens and rhomboid) and medial septum, 100 nL of virus were injected at a rate of 20 nL/min at the following coordinates: entorhinal cortex -4.7 AP, - 4.1 ML, -2.7 DV; ventromedial thalamus -0.7 AP, 0 ML, -3.8 DV; medial septum +0.9 AP, 0 ML, -3.8 DV. For CA1 hippocampus, two injections of 250 nL were performed at a rate of 25 nL/ min (-1.8 AP, -1.6 ML, -1.25 DV and -2.3 AP, -2.4 ML, -1.25 DV). Entorhinal cortex injections mostly targeted the lateral entorhinal cortex. At least two weeks were allowed for Chronos and jRGECO1a expression before recording procedures and chronic hippocampal window, respectively.

### *Ex vivo* patch clamp recordings and optogenetics in adulthood

Patch clamp recordings in adult slices were performed using a SliceScope Pro 1000 rig (Scientifica) equipped with a CCD camera (Hamamatsu Orca-05G). Slices were transferred to a submerged recording chamber and continuously perfused with oxygenated ACSF (3 mL/min) at ∼32°C. Patch recording electrodes (4-6 MΩ resistance) were produced as described above. For current clamp recordings, electrodes were filled with an intracellular solution containing (in mM): 125 K-gluconate, 5 KCl, 4 ATP-Mg, 0.3 GTP-Na_2_, 10 Na_2_-phosphocreatine, 10 HEPES and 0.05% neurobiotin (pH 7.3 and ∼280 mOsm). For voltage clamp experiments, an intracellular solution with the following ingredients (in mM) was used: 152 Cs-methanesulfonate, 4 CsCl, 4 ATP-Mg, 0.3 GTP-Na_2_, 10 Na_2_-phosphocreatine, 10 HEPES (pH 7.3 and ∼280 mOsm). Electrophysiological signals were amplified (Multiclamp 700B), low-pass filtered at 2.9 kHz, digitized at 5-20 kHz and acquired using a Digidata 1440A digitizer and pClamp 10 software (all from Molecular Devices).

Patch clamp recordings were performed from visually identified neurons (GFP+ cells for ebGABAs, random putative GABAergic cells for ctrlGABAs). EbGABAs and ctrlGABAs were sampled from roughly equal numbers from CA1 layers to avoid biases towards specific cell types or inputs. We ensured that no pyramidal cell was included in the ctrlGABA dataset: first, we avoided pyramidal-shaped neurons in the strata pyramidale and oriens; second, we verified in current clamp experiments that all targeted cells (21/21) displayed stereotypical GABA cell firing; third, we verified histologically a subset of ctrlGABAs that were filled with neurobiotin (*n* = 28, see below for details), and all of them exhibited GABA cell anatomical features.

An optoLED system (Cairn Research) consisting of two LEDs was used to visualize fluorescence signals and stimulate Chronos-expressing afferents in CA1. A 470 nm LED coupled to a GFP filter cube was used to visualize GFP-expressing cells and activate Chronos-expressing terminals (3 ms-long light pulses). A white LED coupled to a TRITC/Cy3 filter cube was used to visualize the Chronos/tdTomato-expressing axons. Cells were tested for postsynaptic responses and included in connectivity data only if located in a part of CA1 densely innervated by tdTomato+ axons. Light was delivered using a 40× objective, leading to a light spot size of ∼1 mm, which was able to illuminate all CA1 layers.

Electrical stimulation of the Schaffer collateral and evoked local IPSC experiment were performed by placing a bipolar stimulating electrode made by two twisted nichrome wires in the stratum radiatum of CA1 (between the recorded cell and CA3). The bipolar electrode was connected to a DS2A Isolated Voltage Stimulator (Digitimer Ltd.) delivering 0.2 ms-long stimuli. Postsynaptic current amplitude and paired pulse ratio were assessed by two stimulations separated by 50 ms. Sweeps were separated by 20 s delay to avoid the induction of plasticity and ensure stable responses. Maximum PSCs were determined by delivering increasing stimulation powers and constructing stimulation power *versus* PSC amplitude curves. Since these curves usually saturated, the maximum PSC amplitude was measured from the first PSC of the plateau. In the few cases in which the amplitude did not saturate, the response obtained from the maximum stimulation power was used to calculate the maximum amplitude.

For sPSCs experiments, R_s_ was not compensated because this allowed to monitor more precisely its changes throughout the experiment. For evoked PSC experiments, 60% compensation was applied to the R_s_. Only recordings with R_s_ <30 MΩ were included in the dataset. The cell was discarded if R_s_ changed by more than 20% throughout the protocol. For electrically-evoked and optically-evoked PSCs, PSC onset delays were consistent with monosynaptic responses (3.3 ± 1.6 ms, mean ± standard deviation).

### Analysis of *ex vivo* patch clamp recordings in adulthood

Analysis of intrinsic membrane properties was performed using custom-made MATLAB scripts. The resting membrane potential was estimated by averaging a 60 s current-clamp trace recorded at 0 pA holding current. The input resistance was calculated from the slope of steady-state voltage responses to a series of subthreshold current injections lasting 500 ms (from -50 pA to last sweep with a subthreshold response, 5 or 10 pA step size). The membrane time constant (τ) was estimated from a bi-exponential fit of the voltage response to a -30 pA hyperpolarizing pulse. The membrane capacitance was calculated as the ratio between membrane τ and input resistance. The first spike in response to a juxta-threshold positive current injection was used to determine: the threshold potential (the first point >0.5 in the first derivative), the fast afterhyperpolarization (calculated from the threshold potential), the action potential half-width (the width at half-amplitude between the threshold potential and the peak of the action potential). The rheobase (in pA) was determined as a 50 ms current injection, able to generate a spike in 50% of the cases in 10 trials. A 1 s-long current injection of 2× the rheobase was used to determine: the firing rate, the adaptation index (range 0–1, defined as the ratio between the mean of first three and the last three inter-spike intervals and the burst index (the ratio between the number of spikes in the first 50 ms and the entire depolarizing step). The sag ratio was calculated by injecting a 1 s-long negative current able to hyperpolarize the cell between -100 and -120 mV, and by dividing the amplitude of the hyperpolarization at the end of the current step by the peak of the hyperpolarization at the beginning of the step (sag ratio close to 0 implies high sag, whereas sag ratio close to 1 implies little or no sag). The maximum firing rate was defined as the maximum value reached in a f/I curve with positive current injections from +20 pA to +720 pA (step size: 50 pA).

Analysis of spontaneous postsynaptic currents (sPSCs) was performed with MiniAnalysis (Synaptosoft). PSC parameters were calculated from two minutes recording time. The decay time (10-90%) was calculated from a mono-exponential fit of mean sPSCs. Analysis of evoked postsynaptic current data was performed with Clampfit 10 (Molecular Devices) and MATLAB custom-made scripts. Since membrane and synaptic parameters may be affected by age, for all adult datasets we tested that the distributions of the ages of the mice used did not differ statistically between ctrlGABA and ebGABA. Additionally, for optogenetics experiments we verified that the time of opsin expression did not differ between the groups.

### *In vivo* calcium imaging in adulthood

A chronic cranial window was implanted using previously published procedures^41^. Mice were head-fixed on a non-motorized treadmill allowing self-paced locomotion (adapted from ref. ^42^) All experiments were performed in the dark. No reward was given. After three to five habituation sessions, mice were alert but calm and alternated between periods of moving and resting activity during imaging. The treadmill was made from a 180 cm black velvet seamless belt for the experiments done in the absence of external cue. The movement of the treadmill was monitored using two pairs of LED and photo-sensors that read patterns on a disc coupled to the treadmill wheel, similarly to what previously described^41, 43^.

For all experiments, extra sound, odor, touch, and light were minimized during the imaging session. Imaging was performed with a single beam multiphoton laser scanning system coupled to a microscope (TriM Scope II, Lavision Biotech). The Ti: sapphire excitation laser (Chameleon Ultra II, Coherent) was operated at 1040 nm for jRGECO1a excitation and 920 nm for GFP excitation. Fluorescence emission was acquired using a 16x objective (Nikon, NA 0.8) and split in two detectors (GaSP PMT, H7422-40, Hamamatsu) with bandpass filters of 510/42 nm for GFP and 580/624 nm for GECO. Scanner and PMT were controlled by a commercial software (Imspector, Lavision Biotech). To optimize the signal-to-noise ratio of fluorescence variation, we used a dwell time exposition of 1.85 µs and a spatial resolution of 2 µm/pixel that allowed us to acquire at 9.85 Hz at a field of view of 400 × 400 µm. Locomotion and imaging triggers were synchronously acquired and digitized using a 1440A Digidata (Axon instrument, 2 kHz sampling) and the Axoscope 10 software (Molecular Devices).

### Analysis of *in vivo* calcium imaging data in adulthood

*In vivo* calcium movies were pre-processed using the CaImAn toolbox for MATLAB^44^. First, movies were motion-corrected using a rigid registration method^45^. Then, contours and calcium transients were detected using a constrained non-negative matrix factorization framework allowing denoising and demixing of fluorescence signals^46^. To ensure correct segmentation of somatic calcium activity, the automatic detection was manually refined by adding and removing ROIs using the correlation image based on neighboring pixels. Fluorescence signals were then deconvolved after detrending and extraction of the 20% F/F. An additional manual refinement was carried out with visual inspection of each ROI in the correlation image and the corresponding trace. Unstable or noisy traces were removed because this led to spurious spike inference. A Markov Chain Monte Carlo approach^47^ initialized by the fast OOPSI algorithm^48^ was used to model rise and decay time constants using a second order autoregressive process, which allowed spike inference from the fluorescence traces. A final visual inspection of overlapping calcium traces and spike raster plots was performed to ensure that the spike times reflected the dynamics of the fluorescence traces.

Temporal alignment of the treadmill movement signal and spike raster as well as subsequent analyses were performed using custom-made MATLAB scripts. Locomotion epochs were defined as time periods with deflections in the photo-sensors signal reading the treadmill movement. Rest epochs were defined as periods > 200 ms without treadmill movement. Two methods were used to define cells activated by locomotion. In the first method, peri-stimulus time histograms for the locomotion onset (PSTH^LOC^) were generated (ten seconds window). The mean 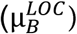 and the standard deviation 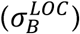 of the baseline firing rate (activity preceding 200 ms before locomotion onset) were used to generate Z-score normalized PSTHs:

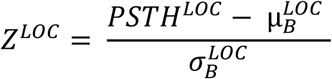

Cells were defined as significantly activated if at least two consecutive bins exceeded a Z-score of 2 in a time window from 200 ms before locomotion onset onwards. This method defined cells activated around the onset of locomotion. In the second method, spike times were circularly shifted to disrupt their temporal relationship with locomotion and rest periods. The ratio between the number of spikes occurring during locomotion and during rest was calculated for each cell of the original and the reshuffled spike matrices. The 95^th^ percentile of the vector containing these ratios for the reshuffled spike times was used as statistical threshold for each cell. Cells were defined as activated by locomotion if their locomotion/rest spike ratio exceeded this threshold. This method defined cells that were broadly more active during locomotion than rest (but not necessarily activated at the onset of locomotion). For each imaging session, a cell was defined as activated by locomotion if it passed at least one of the above tests.

Synchronous calcium events were detected from rest periods binning the spike raster matrix with a 200 ms window. We circularly shifted each spike vector, hence maintaining each cell’s firing rate, but disrupting its temporal relationship to the rest of the population. One thousand surrogate distributions were created. The spikes of each frame for these distributions were computed and the 99^th^ percentile of the resulting “sum of spikes” vector was used as a statistical threshold. Peaks above the threshold that were at least separated by one second were considered SCEs. To define cells activated during SCEs, PSTHs for SCE onset (PSTH^SCE^) were constructed (four seconds window). The mean 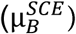 and the standard deviation 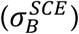 of the baseline firing rate (activity preceding SCEs) were used to generate Z-score normalized PSTHs:

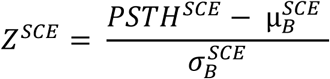

Cells were defined as significantly activated if the bin at zero time lag exceeded a Z-score of 3.

Neuronal assemblies (co-active neurons) were detected using an unsupervised statistical method based on independent component analysis^49^. We chose this method because it allows the analysis of assembly activities over time. Analysis was restricted to rest periods and a 200 ms time binning was used because assembly activations coincident to sharp-wave ripples are known to occur in these conditions^25^. Binned spike counts were z scored to reduce the influence of firing rates. The number of significant co-activation patterns was estimated as the number of principal component variances above a threshold derived from the circularly shifted spike count matrix. Assembly patterns were then extracted with an independent component analysis. To track the activation of cell assemblies over time, a projection matrix was constructed for each pattern from the outer product of its weight vector. To achieve a high temporal resolution, the spike train of each neuron was convolved with a Gaussian kernel (and then z scored).

The activity of each assembly was estimated by projecting the columns of the spike matrix (time bins) onto the axis spanned by the corresponding assembly pattern (principal components). The length of the projection was calculated by taking the inner product between the assembly pattern and a weighted sum of the Z-scored spike counts^49^.

To define cells significantly modulated by assembly activities, we cross-correlated the spike count vector of each cell to the activity vector of each assembly (eight seconds time window). For each cell, we averaged cross-correlograms for different assemblies to define ‘mean cross-correlations to assembly activity’. To calculate a statistical threshold for mean cross-correlations, the same procedure was performed on reshuffled data (circularly shifted spike count matrix) and the 95^th^ percentile of this surrogate distribution was set as threshold for each cell. Mean cross-correlograms with at least two consecutive bins out of the five bins around zero time lag exceeding this threshold were defined as significant.

### Histological processing

Slices containing neurobiotin-filled cells were fixed overnight at 4°C in PFA (4%), rinsed in PBS containing 0.3% Triton X-100 (PBST) and incubated overnight at room temperature in streptavidin -488, -555, -594 or -649 (1:1000 in PBST). Post hoc analysis was performed using a confocal microscope (Leica TCS SP5-X) equipped with emission spectral detection and a tunable laser providing excitation range from 470 to 670 nm. Stacks of optical sections were collected for computer-assisted neuron reconstructions.

Primary antibodies (Table 1) were detected with fluorophore-conjugated secondary antibodies for wide-field epifluorescence and confocal microscopy. After preincubation in 10% normal donkey serum (NDS) in PBST, sections were incubated with a mix of up to three primary antibodies simultaneously diluted in PBST with 1% NDS. The following secondary antibodies were used (all from Jackson Immunoresearch): donkey anti-chicken Alexa 488 (1:1000, 703-545-155), donkey anti-rat Cy3 (1:500, 712-165-150), donkey anti-sheep Dylight 647 (1:250-1:500, 713-495-147), donkey anti-rabbit Dylight 594 (1:500, 711-515-152), donkey anti-rat Alexa 594 (1:500, 705-585-003). For immunostainings with multiple antibodies, an initial negative control was performed by omitting each primary antibody in turn from the staining procedure; in these cases, no positive fluorescence signal was detected. Additionally, each secondary antibody was omitted in turn to confirm its specificity. Epifluorescence images were obtained with a Zeiss AxioImager Z2 microscope coupled to a camera (Zeiss AxioCam MR3) with an HBO lamp associated with 470/40, 525/50, 545/25, 605/70 filter cubes. Confocal images were acquired either with the Leica system described above or with a Zeiss LSM-800 system equipped with emission spectral detection and a tunable laser providing excitation range from 470 to 670 nm.

**Table 1.**
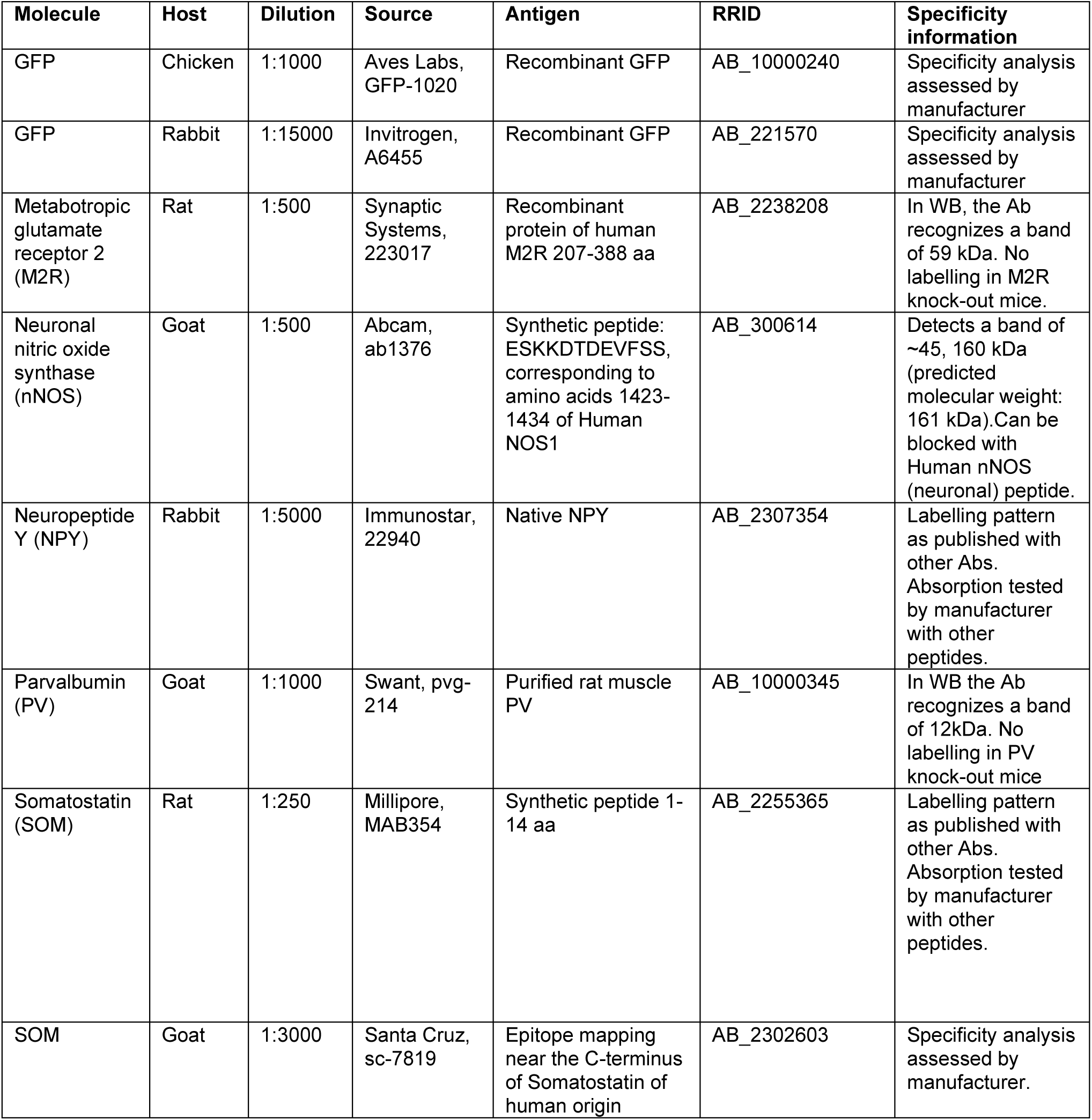
List of primary antibodies.

### Quantification of PV-expressing axon terminals

The innervation of ctrlGABA and ebGABA by PV+ axon terminals was assessed in PFA-fixed sections stained for PV from GAD67-GFP and Dlx1/2(E7.5)-GFP mice, respectively. Confocal stacks centered on the soma and proximal dendrites of GFP+ cells were acquired with a Zeiss LSM-800 microscope at constant resolution (0.065 µm/pixel) and z-step (0.41 µm). Since PV+ axon density varies depending on CA1 layers, ctrlGABA swere sampled to match as much as possible the location of ebGABAs (ctrlGABAs: 17 from stratum oriens, 26 from intra-/peri-stratum pyramidale, 4 from stratum radiatum; ebGABA: 16 from stratum oriens, 14 from intra-/peri-stratum pyramidale, 2 from stratum radiatum). Appositions between PV+ boutons and GFP+ somata or proximal dendrites were counted manually using the cell counter plugin in Fiji (http://fiji.sc). Area and median fluorescence of the PV staining were quantified using first the threshold function to exclude unspecific signal.

### Neurolucida reconstruction and morphometric analysis

Fifty-two neurobiotin-filled neurons (38 filled in developing slices, 14 filled in adult slices) were reconstructed using Neurolucida (MBF Bioscience). Neurons recorded during development underwent morphometric analysis. Examined morphological variables included: dendritic and axonal lengths, dendritic and axonal surfaces.

### Estimate of long-range projecting neurons originating from ebGABA

The proportion of ebGABAs formed by long-range projecting cell types was estimated as follows. We summed: (1) the percentage of ebGABAs formed by SOM+ (but nNOS-) cells in stratum oriens; (2) the percentage of ebGABAs formed by strong nNOS+ cells; (3) the percentage of ebGABAs formed by M2R+ (but nNOS-) cells.

SOM+ cells in stratum oriens are very likely to be projection cells for three reasons. First, most of them are not O-LM cells (very few express PV and we never observed a dense axonal plexus in lacunosum-moleculare in filled cells). Second, most of them are not bistratified cells because very few ebGABAs co-express NPY and PV^50^. Third, the majority of septum-projecting CA1 GABA cells are ebGABAs with the soma in the stratum oriens, and ∼80% of them express SOM^12^.

### Drugs

NBQX disodium salt and SR95531 (gabazine) were purchased from Tocris Biosciences. All the remaining drugs (tamoxifen and compounds to prepare ACSF and intracellular solutions) were purchased from Sigma.

### Statistical analyses

Pairwise comparisons between distributions were performed using the Mann-Whitney U test. Comparisons between multiple groups were performed using the Friedman test with *post hoc* Dunn’s correction. Two-way ANOVAs were performed with Bonferroni *post hoc* correction. Data are expressed as either means ± standard deviations or medians and inter-quartile ranges (IQR). Statistical tests were performed with Graphpad Prism 8 (GraphPad Software, Inc.).

### Data and code availability

Calcium imaging data and PV innervation images will be uploaded to an open-access data repository upon publication of this paper. Custom MATLAB codes to analyze intrinsic membrane properties and *in vivo* calcium imaging data will be uploaded in an open-access Git repository. The remaining data and codes are available from the authors upon request.

## Author Contributions

Conceptualization: R.C. Experimental design: R.C., C.G., M.B., A.B. Data acquisition: C.G., M.B., A.B. and T.To. Software: M.B., D.A., T.Tr. Formal analysis: M.B., C.G., D.A. Technical assistance: T.Tr. Supervision: R.C., A.B. Figures: M.B., C.G. Writing: M.B., R.C.

## Acknowledgements

We thank Loris Cagnacci, Catherine Pauchet-Lopez, François Michel (InMAGIC imaging facility) and INMED’s animal facility technicians for excellent technical support. We are grateful to Valerie Crépel and Alfonso Represa for sharing equipment. We thank Vitor Lopes-Dos Santos for sharing the assembly detection code and Yannick Bollmann and Julien Denis for analytical help. We also thank all Cossart lab members for helpful feedback and discussions.

---

**Supplementary table 1 (related to Figure 1).** Effect of stimulation of ctrlGABAs and ebGABAs on GDP occurrence

| GABA cells (#) | Stimulation frequencies (Hz) | GDP rate | GDP synchronization |  |
| --- | --- | --- | --- | --- |
|  |  | Effect and *P*-value | *P*-value | Delay of evoked GDP after stimulation (s) |
| ebGABA #d16 | 0,1 | ns | < 0.001 | 1.2 |
|  | 0,2 | ns | < 0.05 | 0.4 |
|  | 0,4 | ns | < 0.01 | 1.93 |
| ebGABA #d20 | 0,1 | ns | < 0.05 | 3.2 |
|  | 0,2 | ↓ < 0.01 | ns | na |
|  | 0,4 | ns | < 0.001 | 0.16 |
| ebGABA #d29 | 0,1 | ns | ns | na |
|  | 0,2 | ↑ < 0.05 | ns | na |
|  | 0,4 | ns | < 0.05 | 0.96 |
| ebGABA #d33 | 0,2 | ns | < 0.01 | 3.6 |
|  | 0,4 | ↓ < 0.01 | < 0.05 | 2.16 |
| ebGABA #d39 | 0,1 | ns | ns | na |
|  | 0,2 | ns | < 0.05 | 3.76 |
|  | 0,4 | ns | < 0.05 | 2.16 |
| ebGABA #d53 | 0,1 | ns | < 0.05 | 5.84 |
|  | 0,2 | ns | < 0.001 | 2.48 |
|  | 0,4 | ns | < 0.01 | 0.96 |
| ctrlGABA #d66 | 0,2 | ns | < 0.05 | 3.03 |
|  | 0,4 | ns | ns | na |
| ctrlGABA #d67 | 0,2 | ns | ns | na |
|  | 0,4 | ns | ns | na |
| ctrlGABA #d68 | 0,2 | ns | ns | na |
|  | 0,4 | ns | ns | na |
| ctrlGABA #d69 | 0,2 | ns | ns | na |
|  | 0,4 | ns | ns | na |
| ctrlGABA #d70 | 0,2 | ns | ns | na |
|  | 0,4 | ns | ns | na |
| ctrlGABA #d75 | 0,2 | ns | ns | na |
|  | 0,4 | ns | ns | na |
| ctrlGABA #d80 | 0,2 | ns | ns | na |
|  | 0,4 | ns | ns | na |
| ctrlGABA #d82 | 0,2 | ns | ns | na |
|  | 0,4 | ns | ns | na |

**Supplementary figure 1 (related to Figure 1).**
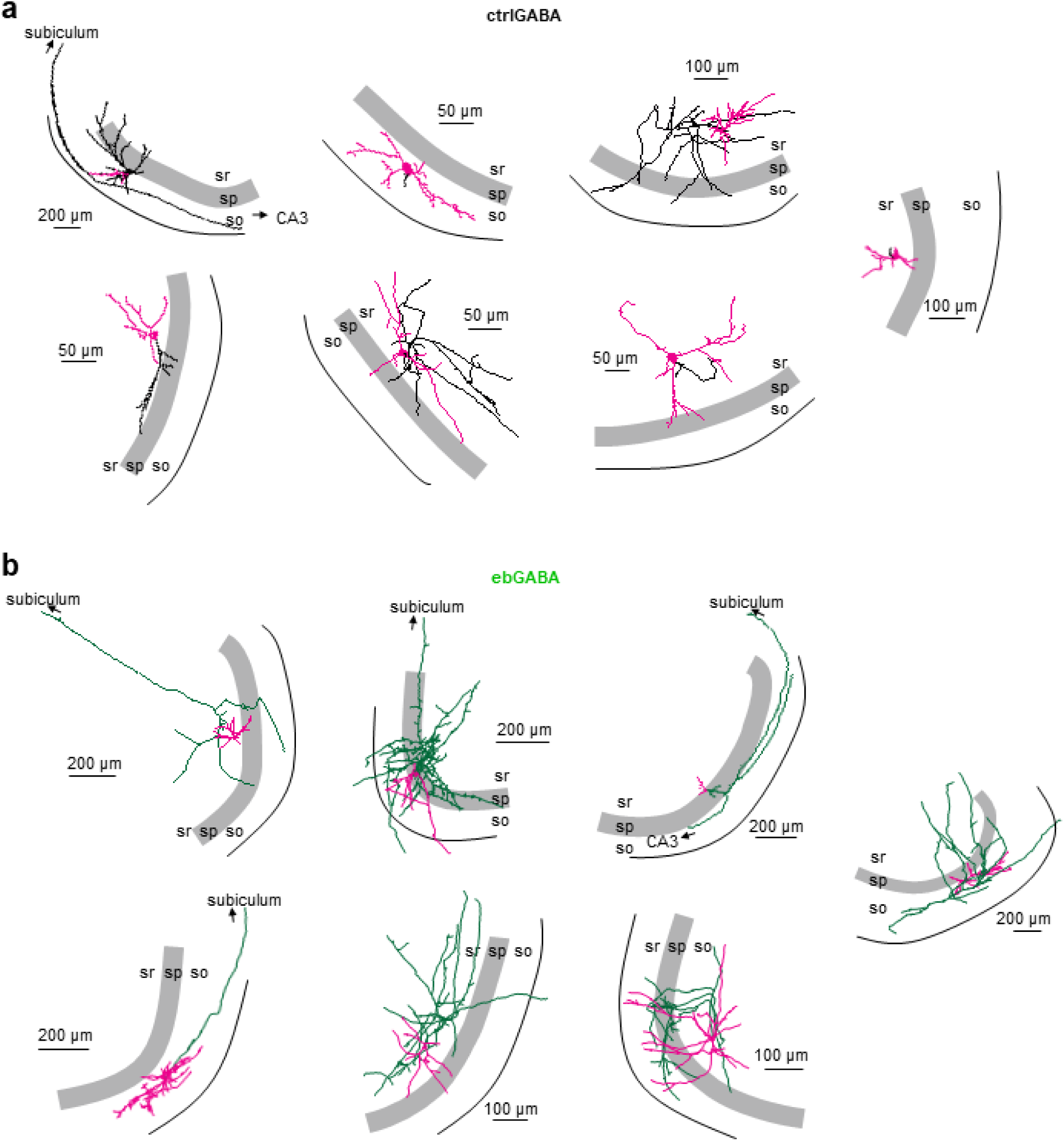
Axonal and dendritic fields of ctrlGABAs and ebGABAs of the developing CA1. Neurolucida reconstructions of seven ctrlGABA and ebGABA cells. Axon is depicted in green for ebGABA and in black for ctrlGABA. Soma and dendrites are colored in magenta. so: stratum oriens; sp: stratum pyramidale; sr: stratum radiatum.

**Supplementary figure 2 (related to Figure 1).**
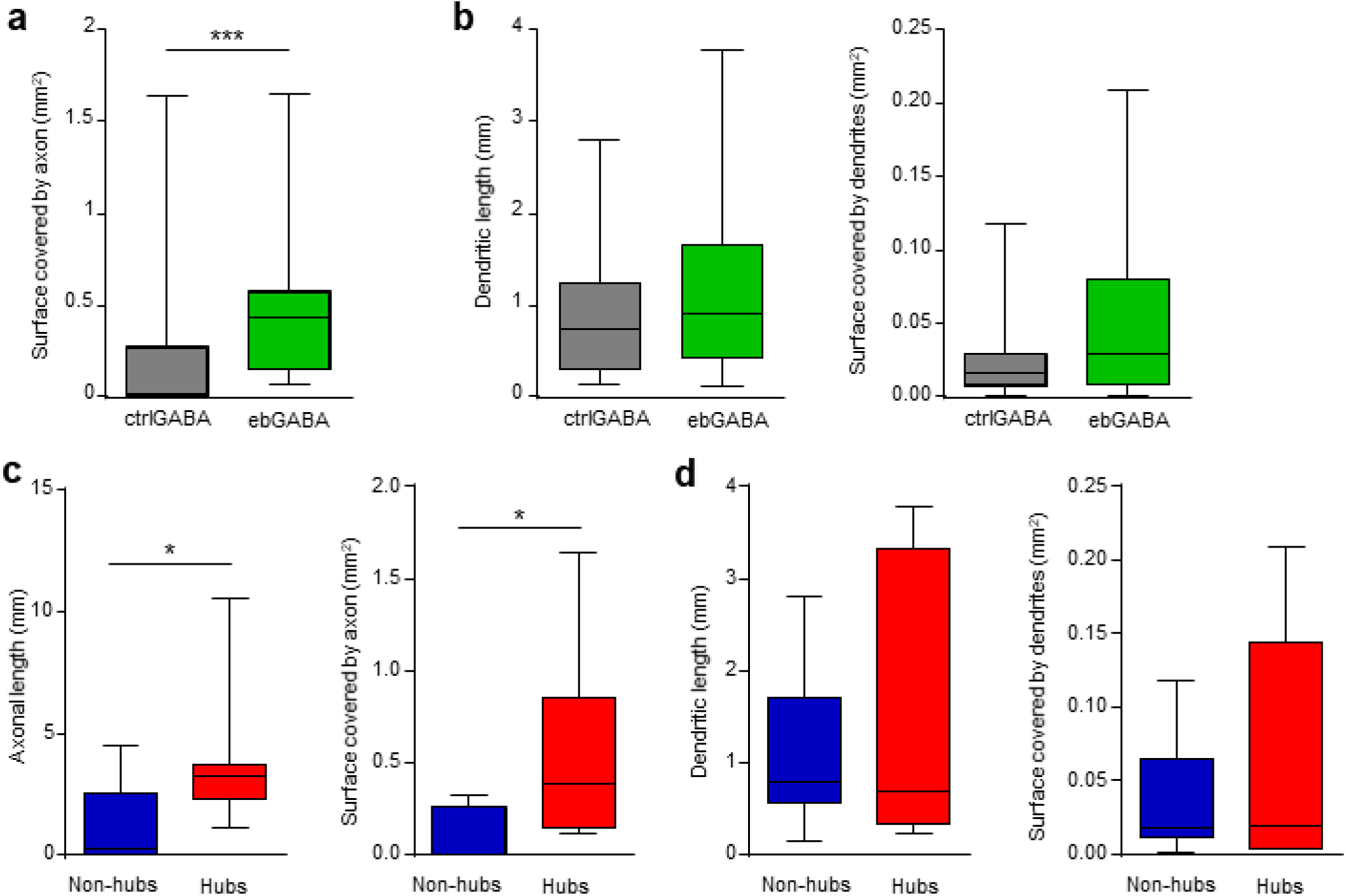
Anatomical features of hub cells in the developing CA1. **(a)** The axons of ebGABAs (*n* = 18) cover a significantly bigger surface than the axons of ctrlGABAs (*n* = 20, *P* = 0.0003, Mann-Whitney U test). **(b)** Dendritic length and surface covered by dendrites do not differ significantly between ctrlGABAs and ebGABAs (*P* = 0.279 and *P* = 0.125, respectively, Mann-Whitney U test). **(c-d)** Cells exerting a hub function in the network (affecting either GDP frequency or GDP synchronization, *n* = 7) display significantly longer axons *(c)* but not dendrites *(d)* than non-hub cells (*n* = 7, axonal length *P* = 0.0379, surface covered by axon *P* = 0.0175, dendritic length *P* = 0.872, surface covered by dendrites *P* = 0.779, Mann-Whitney U test). Data are represented as medians (interquartile ranges). Boxplot whiskers represent minimum and maximum values.

**Supplementary figure 3 (related to Figure 2).**
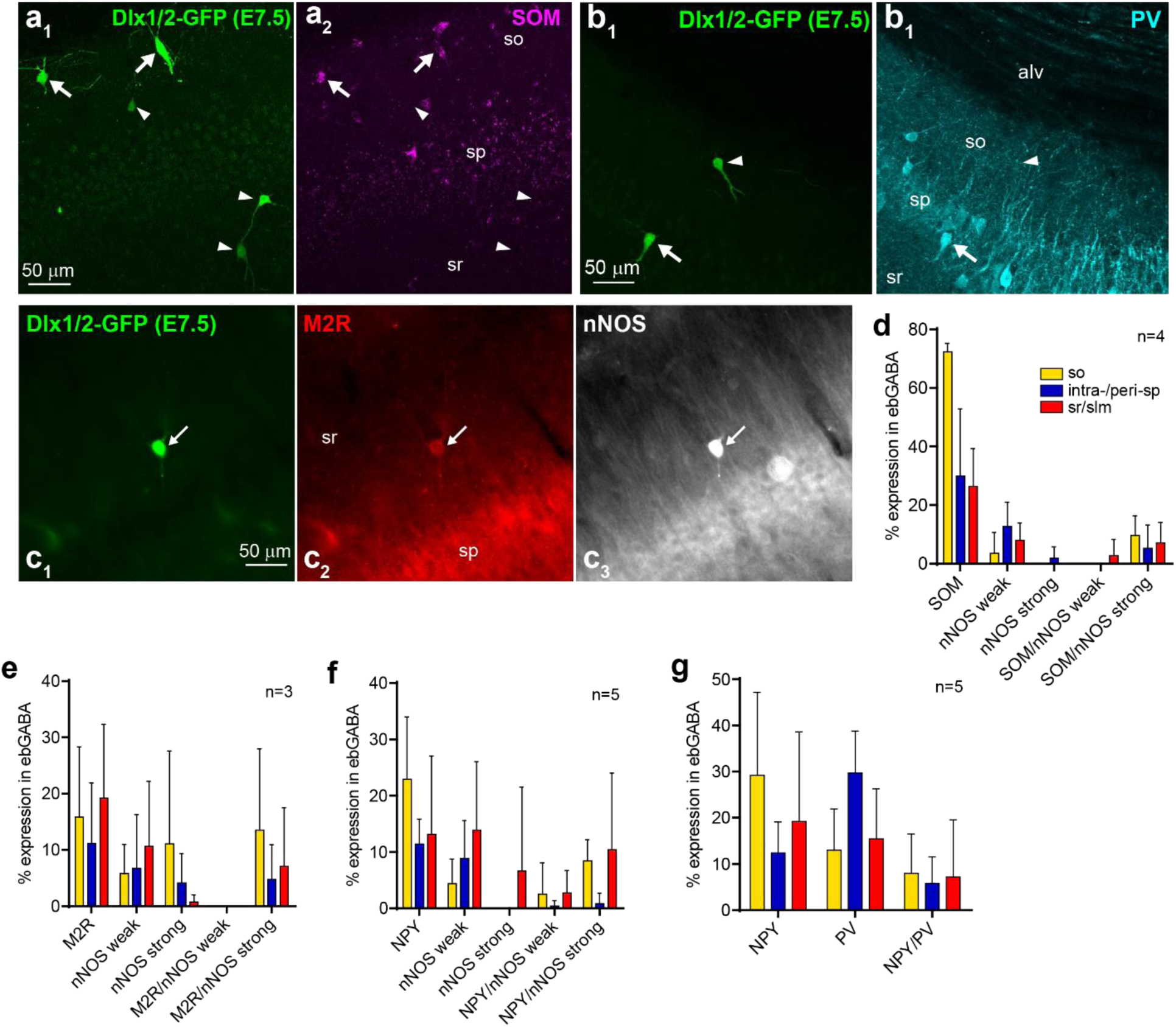
Neurochemical markers expressed by ebGABA in the adult CA1. **(a)** Representative ebGABA cells immunopositive (arrows) or immunonegative (arrowheads) for SOM. Note the higher expression of SOM in stratum oriens. **(b)** One representative ebGABA at the border between the stratum pyramidale and the stratum radiatum expresses PV (arrow), while another ebGABA in stratum oriens is PV immunonegative (arrowhead). **(c)** ebGABA in stratum radiatum expressing both M2R and strong levels of nNOS (arrow). **(d-g)** quantification of the co-expression of two molecular markers in ebGABA. Proportional expression refers to each layer (and not to the whole CA1). Data are represented as means ± SDs. so: stratum oriens; sp: stratum pyramidale; sr: stratum radiatum; slm: stratum lacunosum-moleculare.

**Supplementary figure 4 (related to Figure 2).**
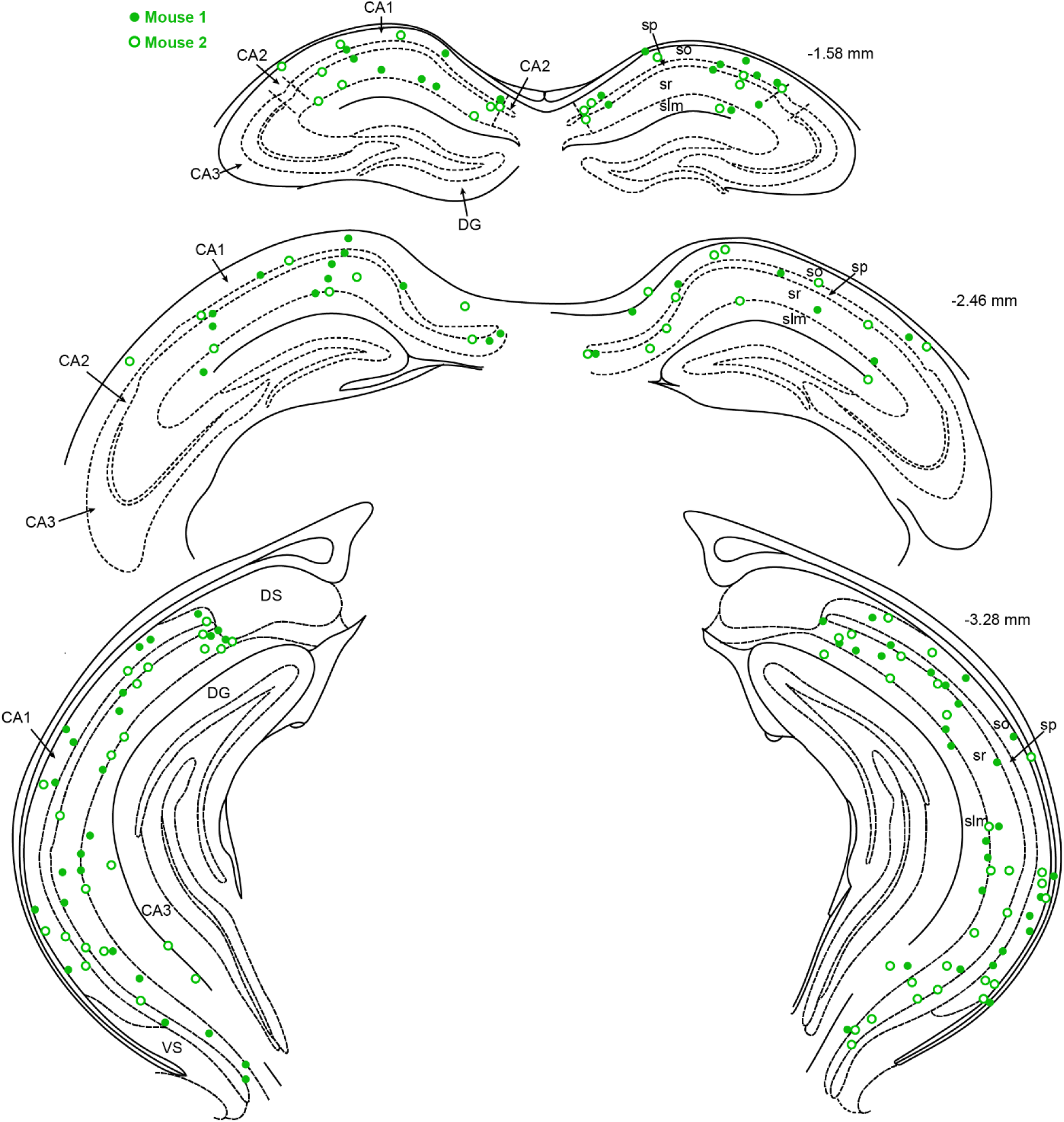
Distribution of ebGABAs in the adult CA1. Position of CA1 ebGABA mapped in two different brains from Dlx1/2(E7.5)-GFP mice at three different rostrocaudal coordinates. At each rostrocaudal level, ebGABA were mapped by collapsing three neighboring 70 µm-thick sections. EbGABAs with somata in CA3, dentate gyrus (DG), dorsal subiculum (DS) and ventral subiculum (VS) were omitted for clarity. so: stratum oriens; sp: stratum pyramidale; sr: stratum radiatum; slm: stratum lacunosum-moleculare.

**Supplementary figure 5 (related to Figure 2).**
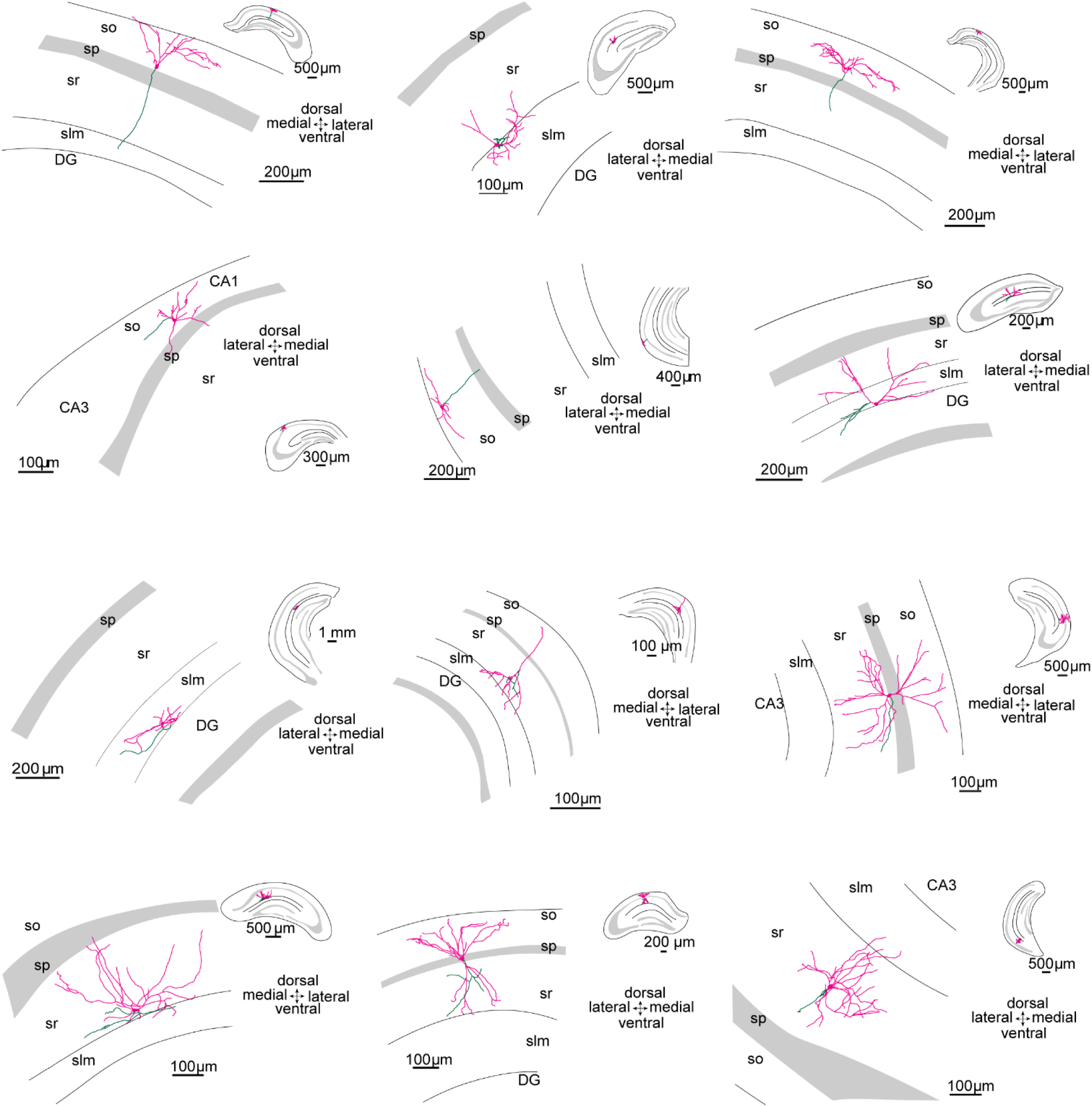
Axonal and dendritic fields of ebGABAs in the adult CA1. Neurolucida reconstructions of neurobiotin-filled ebGABA in the adult CA1. Axon is depicted in green for ebGABA. Soma and dendrites are colored in magenta. Inset: zoomed out representation showing the entire hippocampus to illustrate the rostro-caudal and dorso-ventral position of the neurons. so: stratum oriens; sp: stratum pyramidale; sr: stratum radiatum; slm: stratum lacunosum-moleculare.

**Supplementary table 2 (related to Figure 2).** Intrinsic electrophysiogical properties of ctrlGABAs and ebGABAs in the adult CA1

| Parameter | ctrlGABA n=18 | ebGABA n=16 | P value |
| --- | --- | --- | --- |
| Rheobase (pA) | 49 (27 – 164) | 39 (23 – 82) | 0.165 |
| $V_m$ rest (mV) | -80.0 (-83.0 – -76.8) | -75.2 (-82.1 – -70.2) | 0.125 |
| $R_{in}$ (MΩ) | 320 (123 – 414) | 357 (256 – 561) | 0.327 |
| Membrane $\tau$ (ms) | 44.4 (33.4 – 77.4) | 36.7 (28.6 – 46.1) | 0.108 |
| Capacitance (pF) | 114.4 (88.3 – 674.7) | 89.3 (56.2 – 180.8) | 0.185 |
| Spike threshold (mV) | -42.5 (-44.9 – -38.3) | -42.2 (-45.0 – -35.1) | 0.437 |
| Spike amplitude (mV) | 63.5 (46.3 – 69.7) | 61.1 (56.8 – 66.6) | 0.952 |
| Spike half-width (ms) | 0.7 (0.5 – 0.9) | 0.8 (0.6 – 1.1) | 0.198 |
| AHP<sub>fast</sub> (mV) | 11.0 (8.7 – 15.7) | 14.1 (11.2 – 14.9) | 0.132 |
| **Sag ratio** | **0.472 (0.378 – 0.557)** | **0.590 (0.439 – 0.619)** | **0.028** |
| Firing rate<br>(2x rheobase, Hz) | 21 (13– 74) | 18 (14 – 25) | 0.369 |
| **Maximum firing rate (Hz)** | **123 (60 – 201)** | **74 (55 – 103)** | **0.047** |
| Adaptation index<br>(2x rheobase) | 0.661 (0.552 – 0.746) | 0.741 (0.660 – 0.767) | 0.551 |
| Burst index (2x rheobase) | 0.069 (0.063 – 0.091) | 0.068 (0.056 – 0.077) | 0.449 |
| CV ISI (2x rheobase) | 0.163 (0.131 – 0.266) | 0.181 (0.109 – 0.249) | 0.852 |

Values are expressed as medians (inter-quartile range). Distributions were compared with a Mann-Whitney U test. Significantly different distributions are highlighted in red. $V_m$ rest: resting membrane potential; $R_{in}$ : input resistance; AHP<sub>fast</sub>: fast afterhyperpolarization; CV ISI: coefficient of variation of the inter-spike interval. Red color denotes significantly different parameters

**Supplementary figure 6 (related to Figure 3).**
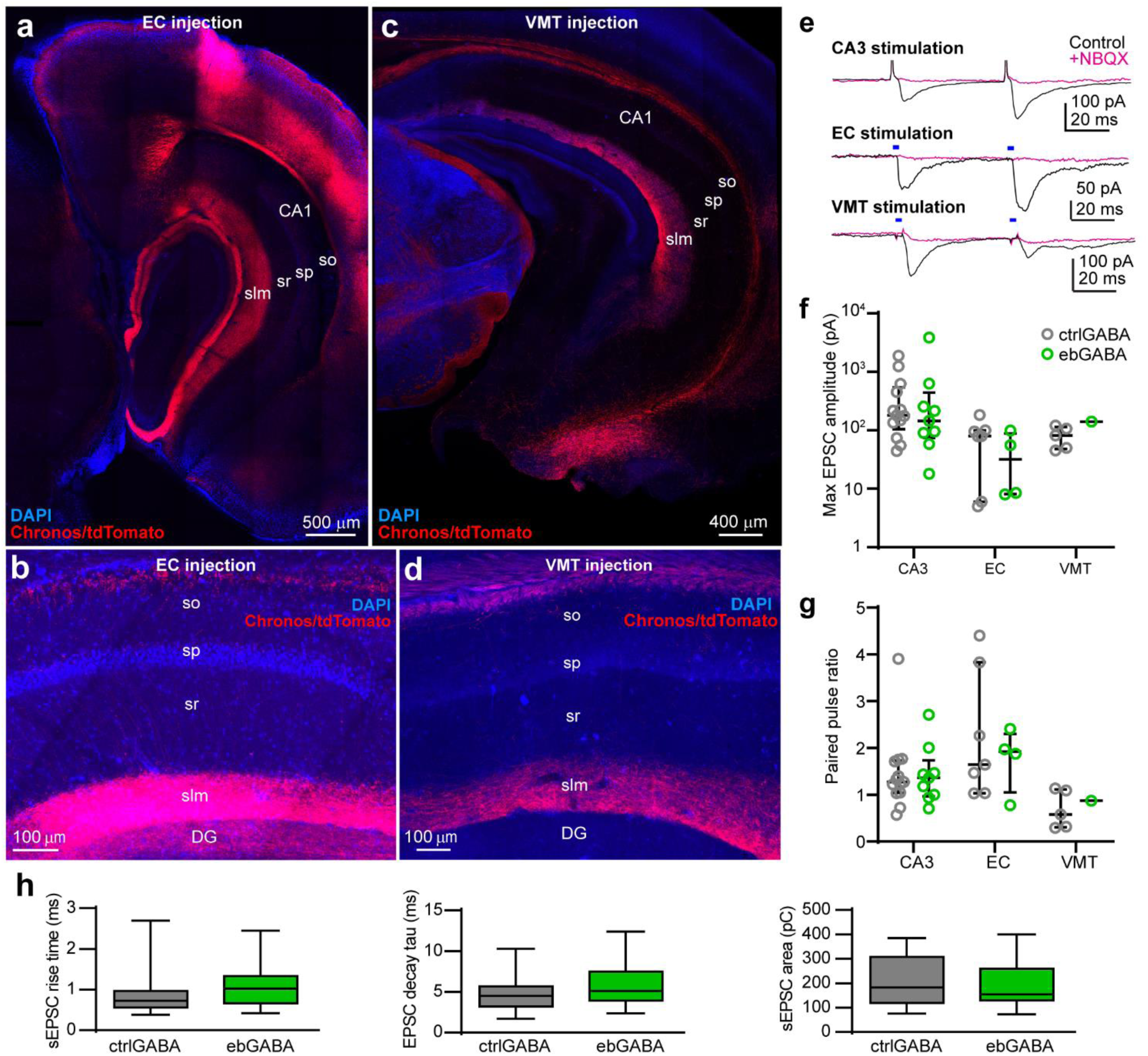
Properties of excitatory inputs to CA1 GABA cells. **(a)** Axons from the entorhinal cortex (EC) innervating the hippocampus. Note that afferents innervating CA1 are densest in the stratum lacunosum-moleculare. **(b)** Zoomed-in view of the entorhinal projection to CA1. **(c)** Axons from the ventromedial thalamus innervating the hippocampus. Note that afferents are restricted to CA1 and, within this region, they are densest in the stratum lacunosum-moleculare. **(d)** Zoomed-in view of the thalamic projection to CA1. **(e)** Three example EPSCs evoked by stimulation of CA3, entorhinal and thalamic axons. All EPSCs are glutamatergic because they are abolished (or greatly reduced) by NBQX (10 μM). **(f-g)** Similar maximum amplitude and paired pulse ratio of EPSCs evoked in ctrlGABAs and ebGABAs from stimulation of CA3 (max amplitude *P* = 0.675, PPR *P* = 0.988), entorhinal (max amplitude *P* = 0.588, PPR *P* = 0.842, Mann-Whitney U tests) or thalamic afferents (no statistics given low number of responses). **(h)** Similar sEPSC kinetics in ctrlGABAs and ebGABAs (rise time *P* = 0.15, decay tau: *P* = 0.8, area *P* = 0.477, Mann-Whitney U tests, both *n* = 18). so: stratum oriens; sp: stratum pyramidale; sr: stratum radiatum; slm: stratum lacunosum-moleculare; DG: dentate gyrus. Data are represented as medians (interquartile ranges). Boxplot whiskers represent minimum and maximum values.

**Supplementary figure 7 (related to Figure 4).**
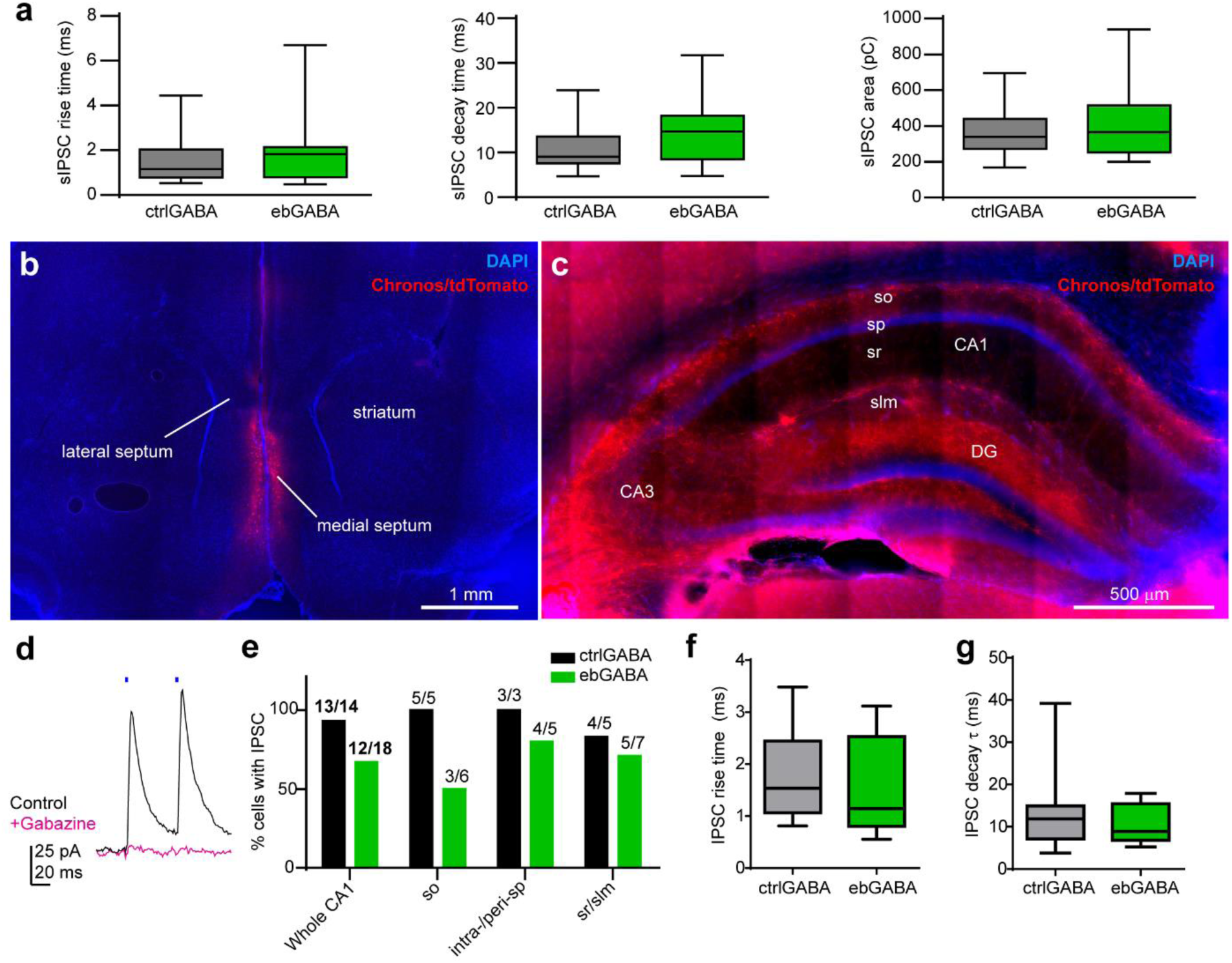
Properties of inhibitory inputs onto CA1 GABA cells. **(a)** The kinetics of sIPSCs do not differ statistically between ctrlGABAs and ebGABAs (rise time: *P* = 0.516, decay tau: *P* = 0.142, area: *P* = 0.6, Mann-Whitney U tests, both *n* = 18). **(b)** expression of Chronos/tdTomato in the medial septum of a representative mouse. **(c)** axons from medial septal neurons innervating the hippocampus. Note that within CA1 afferents innervate all layers but are particularly dense in stratum oriens and at the border between stratum radiatum and stratum lacunosum moleculare. **(d)** IPSC evoked by optical stimulation of medial septal afferents is blocked by gabazine (SR95531, 10 μM). **(e)** Proportion of CA1 ctrlGABA and ebGABA cells receiving GABAergic inputs from the medial septum. No significant difference in the proportions for the whole CA1 (*P* = 0.104, Fisher’s exact test). **(f-g)** Similar septal IPSC rise time (*P* = 0.43) and decay time (*P* = 0.683, Mann Whitney U tests) in ctrlGABAs (*n* = 13) and ebGABAs (*n* = 12). so: stratum oriens; sp: stratum pyramidale; sr: stratum radiatum; slm: stratum lacunosum-moleculare; DG: dentate gyrus. Distributions are represented as medians (interquartile ranges). Boxplot whiskers represent minimum and maximum values.

**Supplementary figure 8 (related to Figure 4).**
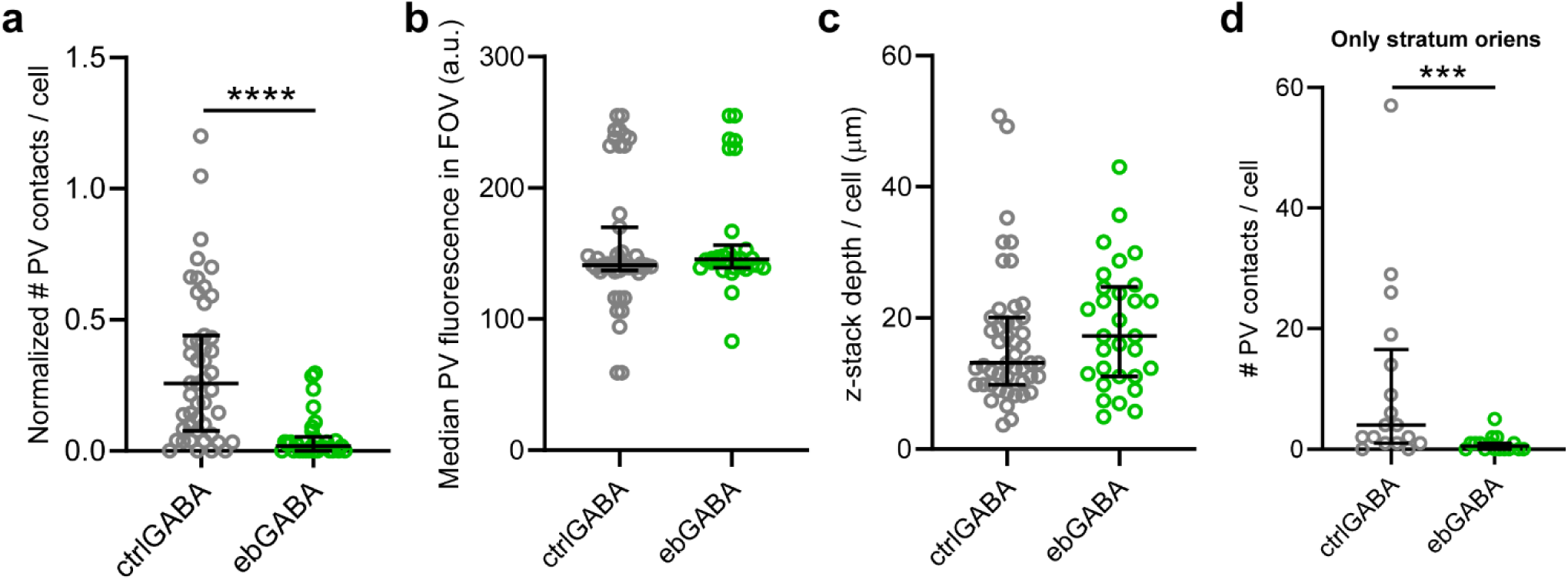
Low amount of PV contacts onto ebGABAs is not due to uneven sampling or staining quality. **(a)** Difference in the amount of PV+ boutons between ctrlGABAs (*n* = 47) and ebGABAs (*n* = 30) persists even when the number of contacts is normalized by the number of optical sections scanned for each cell (*P* < 0.0001, Mann-Whitney U test). **(b)** Similar median fluorescence of the PV staining for examined ctrlGABAs and ebGABAs cells (*P* = 0.275, Mann-Whitney U test). **(c)** No difference in the z-stack depth for imaged ctrlGABAs and ebGABAs (*P* = 0.23, Mann-Whitney U test). **(d)** Difference in the amount of PV+ boutons between ctrlGABAs and ebGABAs persists even when analysis is restricted to cells with the soma in the stratum oriens (*P* = 0.0008, Mann-Whitney U test). *** p < 0.001. **** p < 0.0001. Horizontal lines represent quartiles.

**Supplementary figure 9 (related to Figure 5).**
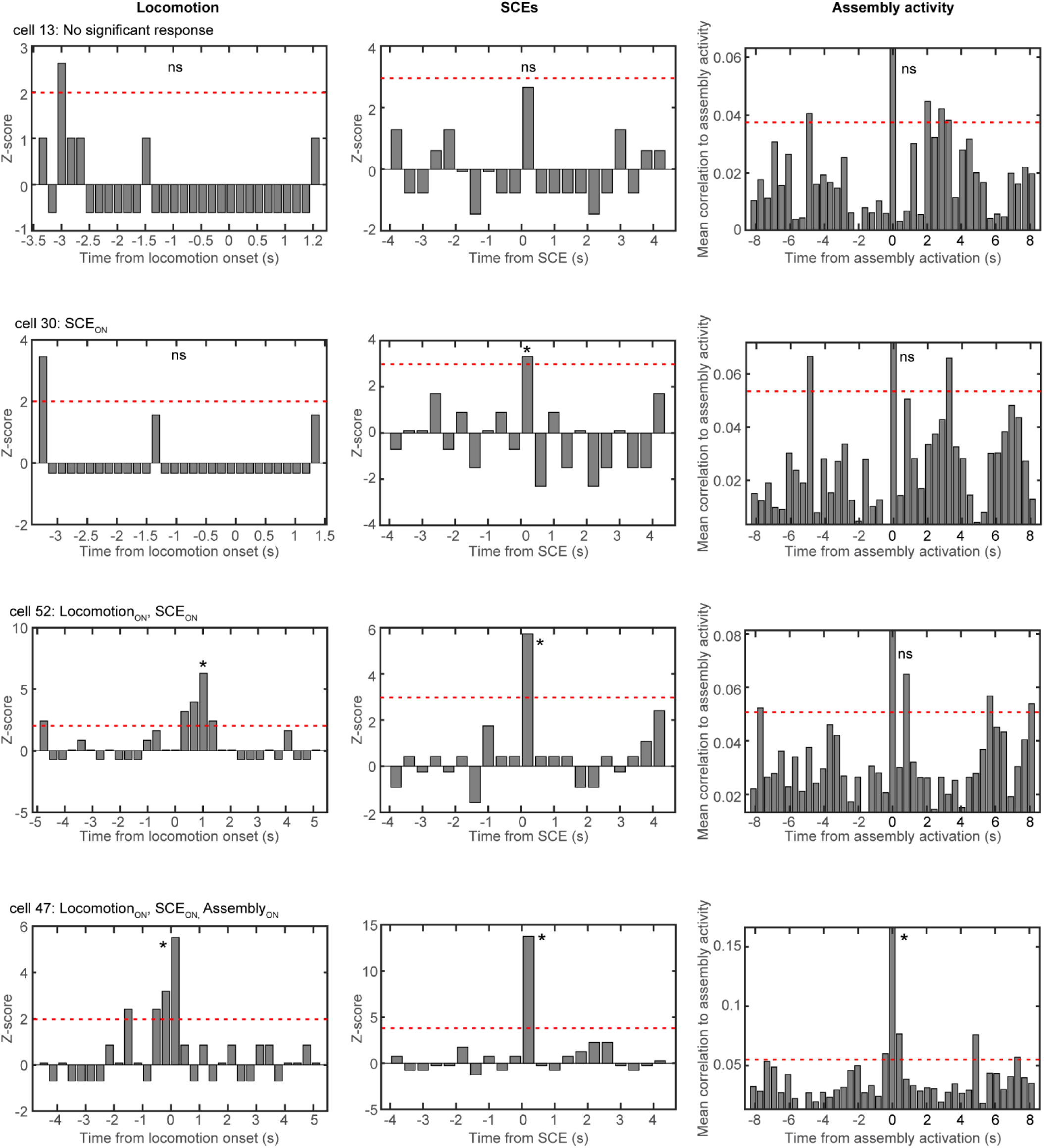
Diversity of single cell responses to locomotion and network events in the adult CA1. Activity of four cells from mouse 2 in relation to locomotion (left), SCEs (middle) and assembly activity (right). Red dashed lines depict statistical thresholds (see Methods for details).

